# ArchLips: A comprehensive *in silico* database for high-throughput identification of archaeal lipids

**DOI:** 10.1101/2025.05.05.652033

**Authors:** Fengfeng Zheng, Wenyong Yao, Wei He, Wan Zhang, Yufei Chen, Huahui Chen, Zhirui Zeng, Xiao-Lei Liu, Su Ding, Yanhong Zheng, Linan Huang, Yuanqing Zhu, Chuanlun Zhang

## Abstract

Archaeal membrane lipids are markedly distinct from those in bacteria and eukaryotes, serving as unique biomarkers for unraveling their ecological and biogeochemical roles. Recent advancements in lipidomics have facilitated detailed cellular-level characterizations of lipid compounds. However, the lack of a comprehensive and dedicated database has severely limited large-scale, high-throughput investigations of archaeal lipids. We present ArchLips, a comprehensive database containing 218,868 *in silico* molecular structures and tandem mass spectra of 199,248 corresponding archaeal lipids. ArchLips enables the rapid and accurate identification of characterized archaeal lipid compounds from both pure cultures and environmental samples, serving as a transformative tool for enhancing our understanding of archaeal diversity and their ecological and evolutionary significance within global ecosystems.

---

Archaea are widely recognized as a unique branch on the tree of life alongside bacteria and eukaryotes and among the earliest microbial forms on Earth^1–6^. Advances in culture-independent high-throughput sequencing and bioinformatic approaches have catalyzed the discovery of a large number of novel and metabolically versatile archaeal species, driving burgeoning research into the diversity, ecological functions, roles in biogeochemical processes, evolutionary history, and biotechnological applications of these organisms. To date, more than 30 archaeal phyla grouped into 4 superphyla (Euryarchaeota, DPANN, TACK, and Asgardarchaeota) have been identified across a wide range of environments, from extreme habitats to diverse mesophilic ecological niches, host-associated microbiomes, and mammalian guts^7–14^. Notably, the recent discovery of the Asgard archaeal superphylum has revealed a close phylogenomic relationship with Eukarya, reigniting debates on origins of eukaryotes and underlining the crucial role of archaea in early evolution of life^8, 15–17^. These milestone findings have prompted intensified research efforts to further elucidate the biological and ecological impacts of these enigmatic microorganisms.

One of the most intrinsic differences between archaea and bacteria/eukaryotes is the structure of their cell membrane lipids. The substantial differences in the stereochemistry of the glycerol backbone (glycerol-1-phosphate in archaea vs. glycerol-3-phosphate in bacteria), the hydrocarbon chains (isoprenoids in archaea vs. fatty acyls in bacteria and eukaryotes), and the nature of the ether/ester bonds linking these components have raised “lipid divide” questions regarding the evolutionary origins and divergence of microbial lipid biosynthesis pathways^6, 18–22^.

The archaeal cell membrane contains monolayer or bilayer lipids consisting of isoprenoid glycerol dialkyl glycerol tetraethers (e.g., iGDGTs) and/or glycerol dialkyl diethers (e.g., archaeol) as core lipids, which can be modified by methylation, cyclization, cross-linking, hydroxylation and unsaturation to increase the structural diversity (Fig. S1). The second level of structural diversity in archaeal lipids arises from the head groups attached to the core lipid structures. These head groups often contain various sugar moieties (glycolipids) and/or phosphate group (glycerophospholipids), which can be further modified with polar molecules such as choline (PC), ethanolamine (PE), serine (PS), thereby forming amphipathic intact polar membrane lipids in archaeal cells (IPLs, Fig. S1). In addition, archaeal cardiolipin analogues have been identified and function as fundamental membrane components in some lineages of archaea^23, 24^. Other non-membrane-forming lipids like quinone or carotenoid have also been identified and involved in electron transfer in some archaea^25^.

These unique lipids not only help archaea survive in diverse environmental conditions but also leave behind as molecular “fingerprints” in sediments and rocks, offering clues about paleoclimates and ecosystems. Yet, this versatility of archaeal lipids or lipidome has only been utilized in a very small proportion for paleoclimate proxy studies such as the TEX_86_ (TetraEther indeX of tetraethers consisting of 86 carbon atoms) proxy for paleotemperature and the methane index (MI) proxy for ancient marine hydrate identification^26, 27^. Increasing evidence shows that archaeal lipids can also reflect paleo ocean redox changes and nutritional status, which are more related to the physiology and biochemistry of archaea^28–30^. Thus, great promises lie ahead with more archaeal lipids being used in paleo-climate as well as paleo-ecosystem studies^31,32^.

Lipidomics is an emerging field in lipid biogeochemistry, both ultra-performance liquid chromatography (UPLC) coupled and shotgun mass spectrometry (MS) approaches have provided unprecedented resolution (MS) for lipid detection and have become benchmark methods for lipidomic research. The untargeted MS approach enables the simultaneous detection of hundreds to thousands of diverse lipid compounds and continues to generate expansive MS-based lipidomic datasets that require high-throughput data processing, particularly for lipid identification. Lipid identification remains a major bottleneck in MS-based lipidomic research, which often relying on manual interpretation or matching experimental data, such as accurate mass and tandem mass (MS^2^) spectra against standard or *in silico* spectral libraries^33, 34^. Due to the limited availability of authentic lipid standards, especially for archaeal lipids, the *in silico* lipid databases have become widely used and show great promise for compound annotation in untargeted MS-based lipidomics^35, 36^. Algorithms such as rule-based modelling or machine learning approaches have been employed to generate *in silico* mass spectra. Incorporating these spectral libraries into extant cheminformatic tools enabled the development of automated and effective methods for MS data interpretation, and expanded coverage of lipids species^34, 37–40^ that would otherwise be impossible for manual annotation.

The advances in lipidomic research have inspired us to build the first comprehensive *in silico* archaeal lipid database, named ArchLips (spreading the word about archaeal lipids), specifically designed for detailed fingerprinting of archaeal lipid structures and compositions. This database covered an extraordinary structural diversity of archaeal lipids, many of which have been only sparsely collected into existing lipid databases (Table 1). Our full library consisted of 199,248 archaeal lipids containing *in silico* MS^2^ mass spectra and can be integrated with bioinformatic tools for high-throughput compound annotation (Fig. S2). The application of the ArchLips database to pure and environmental samples for lipid identification should provide an invaluable resource for promoting archaeal lipidomic research. It will support the discovery of novel lipid biomarkers, expand the scope of archaeal lipids in paleo-ecosystem studies, and help elucidate archaea-eukaryote evolution pathways involving lipid membrane transition.

**Table 1.** Archaeal lipids in the public database of LipidMAPs, PubChem and LipidBank. Please note that the removal of replicates was performed based on the formula.

|  | LipidMAPs | PubChem-Compounds | PubChem-Substances | LipidBank |
| --- | --- | --- | --- | --- |
| Total compounds | 48,548 | 118,564,406 | 319,893,398 | 7009 |
| Archaeal lipid | 26 | 68 | 252 | 232 |
| Archaeal lipid:<br>replicates removed | 26 | 47 | 53 | 145 |

## RESULTS

### Construction of the general structure library

This study began with the compilation of a structure database of archaeal lipids from all available sources, including public databases such as LipidMAPS (https://lipidmaps.org/), PubChem (https://pubchem.ncbi.nlm.nih.gov/), and LipidBank (https://lipidbank.jp/). Significant effort was devoted to collecting publicly available lipid structures through meticulous literature research (e.g., Ref. ^23, 41–54^). We then constructed the structure library that constitutes all currently known archaeal lipids, including their isomers, homologs and derivatives (Table 2 and Fig. S1). This resulted in a total of 218,868 *in silico* molecules in distinct structures (Fig. 1) generated using MarvinSketch, ChemAxon Reactor software and the RDKit package in python, which were stored in SDF (MDL molfile) format.

**Figure 1.**
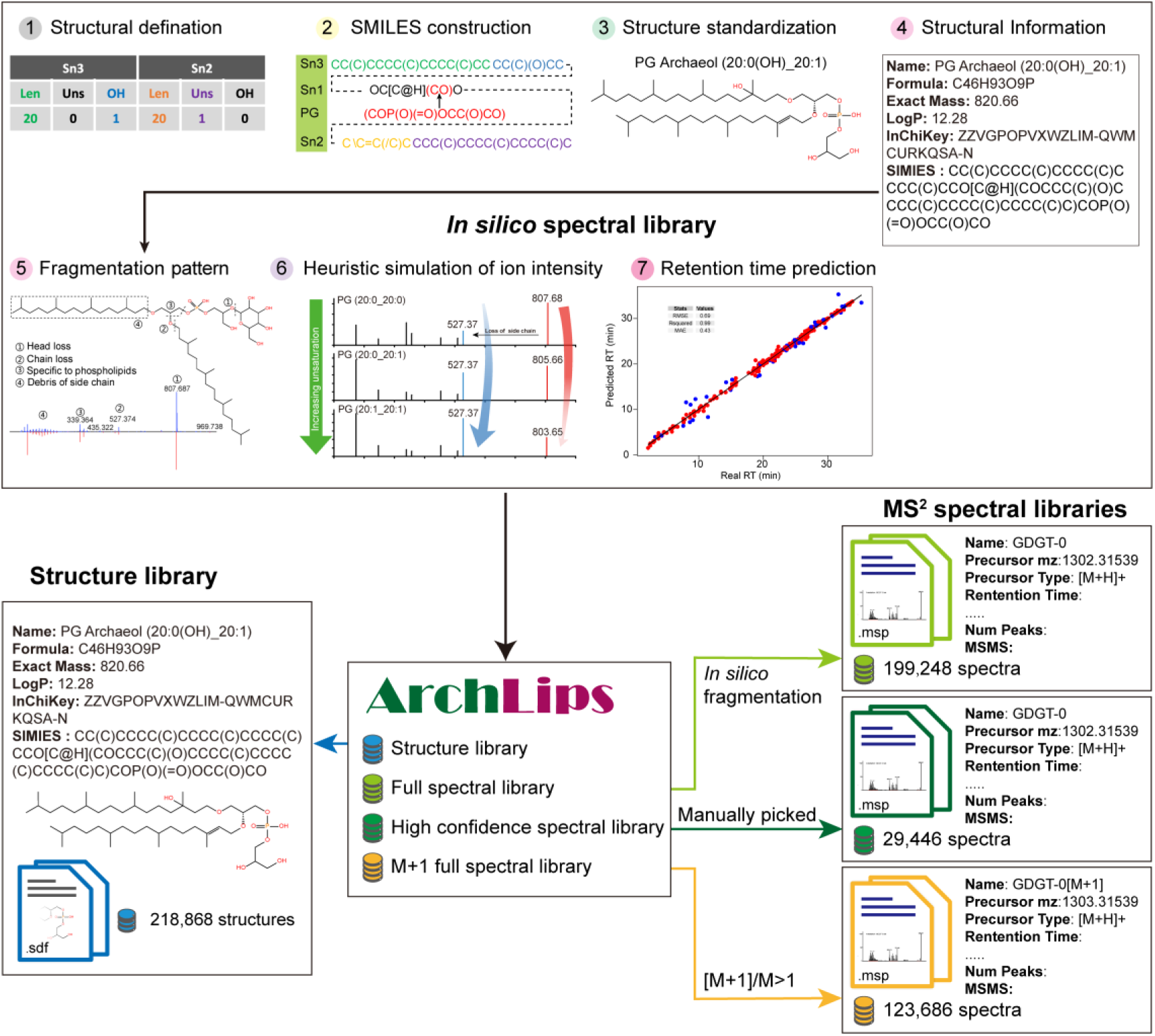
Generation of ArchLips database. The *in silico* structure library was compiled following steps 1-4 using a combinatorial enumeration method based on structures reported in the literature. The in silico MS^2^ spectra were curated according to steps 5-7 based on the reported fragmentation pattern of archaeal lipids, and the ion intensity was calculated in a heuristic way. The ArchLips database contains four libraries: an *in silico* structure library, a full spectral library, a filtered high-confidence library and a library containing the M+1 spectra.

**Table 2.**
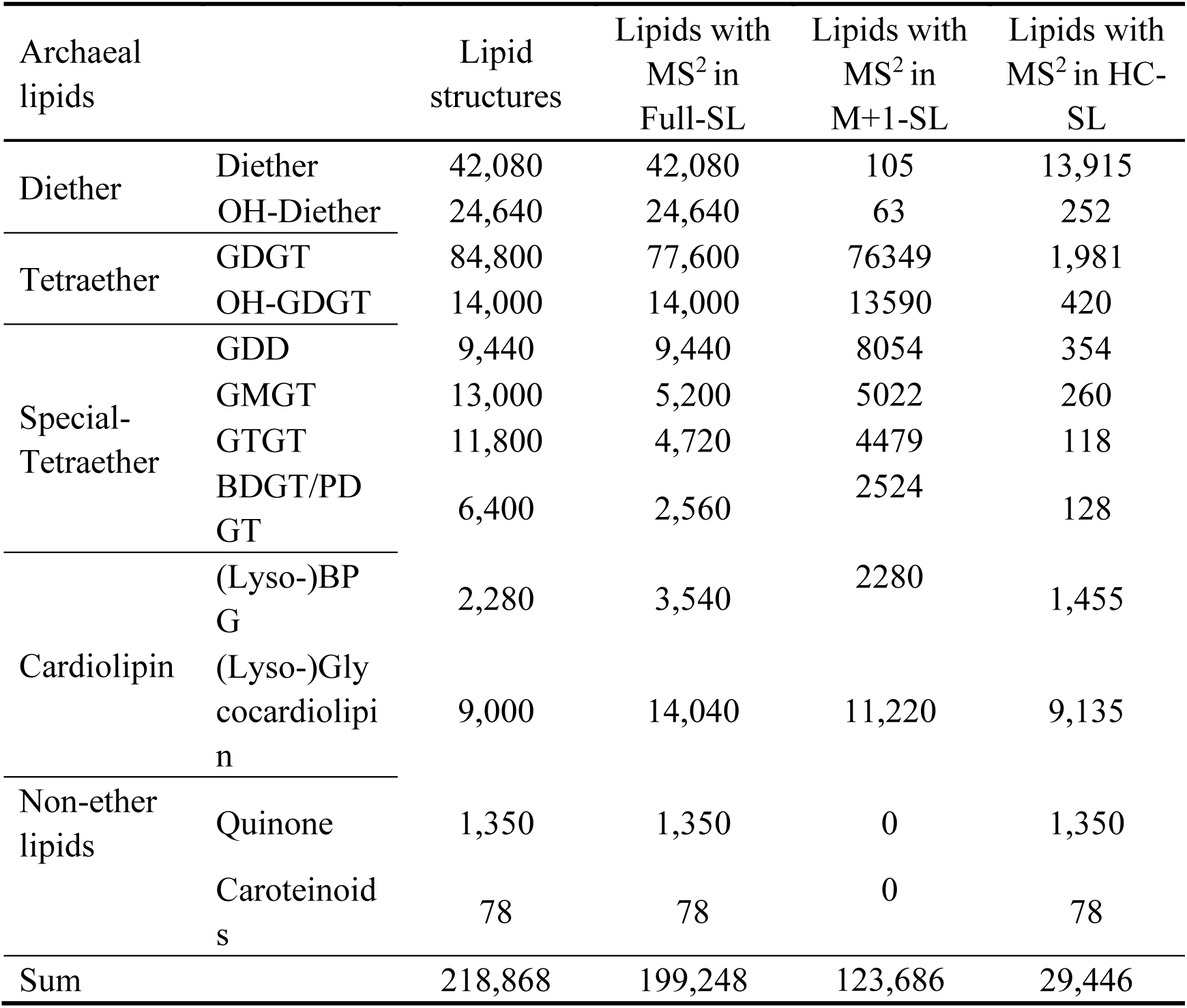
Number of lipid compounds and tandem mass spectra in the ArchLips database. Please refer to Figure S1 for full names and structures of lipids.

The structure library also included basic chemical information such as molecular formula, exact mass, SMILES, Inchi, InchiKey, and logP. The nomenclature of archaeal diether lipids followed that of the LipidMAPS at the molecular level such as PG Archaeol (20:0_20:0) and 2Gly-Archaeol(20:0_20:0). Traditional names of certain compounds, such as TGD-PA, S-DGD-PA were retained for consistency. For tetraether lipids, traditional names were preserved, while regioisomer compounds were labeled using suffixes such as “a,” “b,” and “c” (e.g., GDGT-4a, GDGT-4b, and GDGT-4c to denote isomers of GDGTs with four cyclopentane rings in different configurations)^43, 44^.

To avoid a combinatorial explosion of generated structures^35^, we defined structural boundaries for archaeal lipids. For instance, the isoprenoid carbon chain lengths in diether lipids were limited to C20–C30, with a maximum of two additional methyl or hydroxyl groups occurring at the C3 position of the carbon chains. The library also included macrocyclic archaeol containing 0–4 rings and unsaturated archaeol with up to 12 double bonds. However, stereoisomers and regioisomers such as variations in double bond position or sn2/sn3 positions of the isoprenoid carbon chains, were not considered in this database.

Archaeal GDGTs containing a maximum of 8 cyclopentane moieties served as the primary structures for further generation of tetraether lipids of possible structures constrained by the boundary condition mentioned above. The resulting diverse derivatives were refined to include methylated GDGTs (Me-GDGTs; 1–3 additional methyl groups) and hydroxylated GDGTs (OH-GDGTs; 1–2 hydroxyl groups), GDGTs with cyclohexane rings (S-GDGTs; 1–2 rings), unsaturated GDGTs (1–8 double bounds) and H-shaped configurations (GMGTs). Selected regioisomers, e.g., the regioisomers from the cyclopentyl ring distributions between etherified chains were defined under specific chromatographic conditions^43, 55^. Other stereoisomers such as tetraether lipids with parallel glycerol backbones were excluded from the database as they lack definitive chromatographic conditions to separate them ^56^.

The MDL molfile structures of archaeal intact polar lipids were generated on the foundation of the core lipids described above using a total of 79 head groups that have been reported in the literature and database (Fig. S1 and Table S2). Because archaeal tetraether lipids possess two terminal glycerol units, polar head groups were allowed to attach to both units^41, 57, 58^, with one example being Gly-(PGly)-GDGT (Glycosyl-GDGT-phosphatidylglycosyl) (Table S2).

### Construction of the ArchLips spectral library

We used LipidBlast as a foundational template for developing an *in silico* spectral library of archaeal lipids because it is easy to operate and formulate new structures^35, 36^. Building upon the template we integrated all structural information into a dedicated Excel file to improve data organization, optimize MS^2^ calculations, and streamline library exportation. Specifically, the MS^2^ data was organized into multiple Excel sheets, with each sheet corresponding to a lipid class with different core lipids that allow optimized MS^2^ calculations; the library exportation program was rewritten in Python, employing multiprocessing to efficiently produce spectral library files.

The customized ArchLips spectral library comprised three core components: (1) basic chemical information, e.g exact mass and a defined formula, (2) diagnostic fragment ion *m/z* values, and (3) the relative signal intensities of these fragments. [M+H]⁺ and [M+NH₄]^+^, two primary adducts observed in positive ionization mode, were incorporated into exact mass to construct the *m*/*z* of precursor ions.

While the theoretical basic chemical properties and the *m*/*z* of precursor ions were calculated directly from structural input, the fragment *m/z* and relative intensity served as critical parameters for spectral matching accuracy. These latter two metrics were systematically generated through the following methodology:

Firstly, the *m*/*z* of fragment ion and the relative intensity were intrinsically determined by the stability of chemical bonds for a specific class of lipids and experimentally by the collision energy level used during collision-induced dissociation (CID). We examined the optimal collision energy using 18 archaeal lipids under different levels (Fig. S3). Based on the observed positive correlation between optimal collision energy and *m*/*z* (Fig. S3B), we implemented a ramped collision energy approach: low-mass ions (start at *m*/*z* 100) were fragmented from 10–15 V, while high-mass ions (end at *m*/*z* 2000) an elevated energies from 55–60 V was used to ensure sufficient bond cleavage (Fig. S4). This strategy, tailored to the molecular weight-dependent fragmentation behavior of archaeal lipids, maximized the yield of diagnostic fragment ions and enabled systematic acquisition of high-quality MS^2^ spectra for database construction.

Secondly, the fragmentation pattern of archaeal lipids was delineated into three distinct regions (Regions I, II, and III) as summarized in Ref. ^45^ (Fig. 2). Region I was associated with the neutral loss of small molecular moieties occurring for almost all archaeal lipids and the sequential loss of polar headgroups in intact polar lipids. These losses of small molecular moieties typically include the sequential loss of H2O, C3H4O, and C3H6O2, resulting in product ions of e.g., [M+H-18]^+^, [M+H-56]^+^, and [M+H-74]^+^, respectively (Fig. 2A).

**Figure 2.**
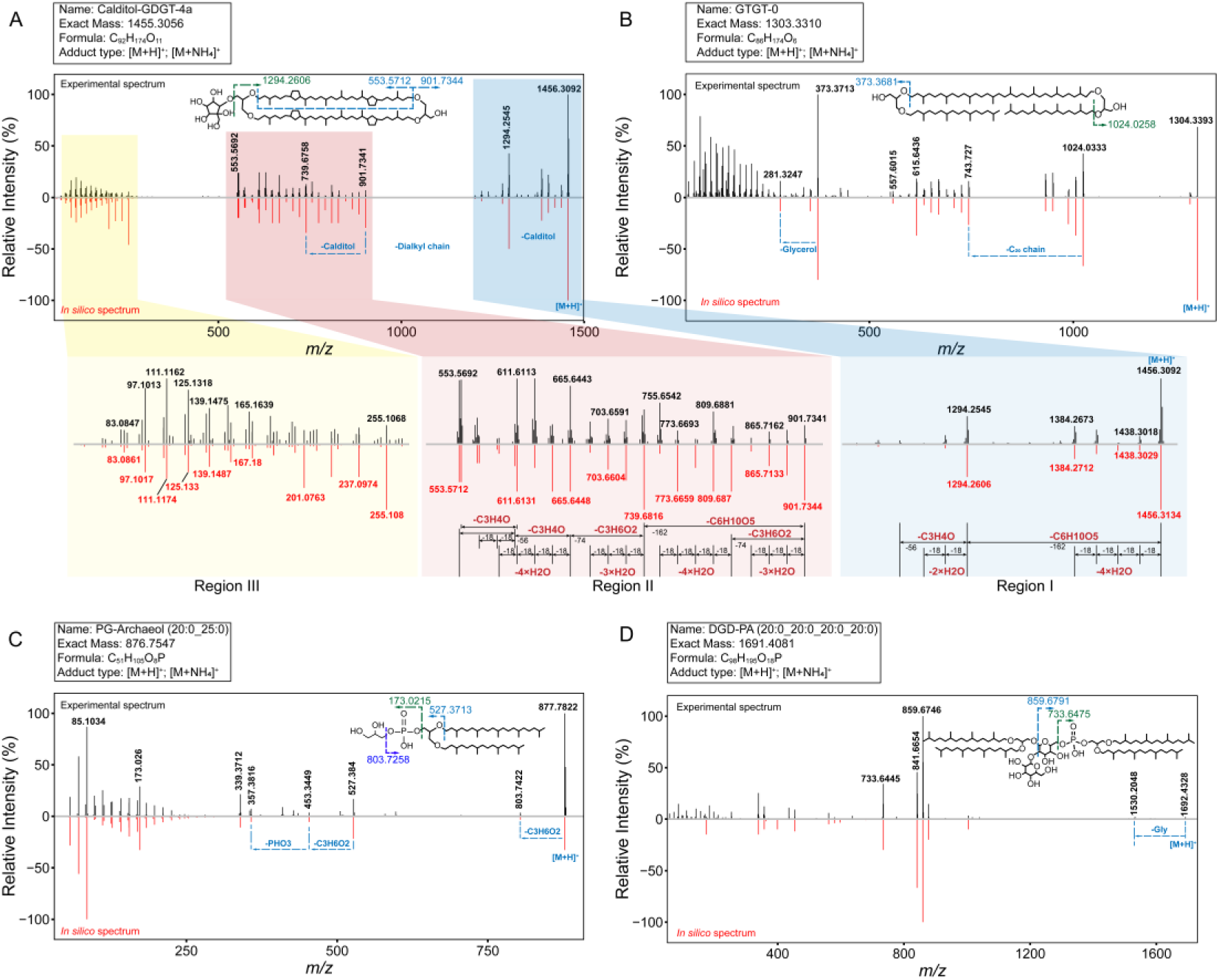
MS^2^ spectra and fragmentation pattern of four typical archaeal lipids. A) MS^2^ spectrum of Calditiol-GDGT-4a with the fragmentation pattern highlighted with three regions (I, II, and III). Region I refers to the product ions from the neutral loss of small molecules, Region II contains the characteristic ions of archaeal lipids that concern the neutral loss of one diphytanyl chain and subsequent loss of small molecules, and Region III involves the product ions from the further cleavage of the biphenyl or phytanyl chains. B) GTGT-0, an intermediate compound during the biosynthesis of archaeal tetraether lipids. C) PG-Archaeol (20:0_25:0), an intact polar lipid with a phosphoglycerol headgroup. D) DGD-PA (20:0_20:0_20:0_20:0), an archaeal cardiolipin analogue. The experimental spectrum is shown at the top (black) and the *in silico* spectrum at the bottom (red).

Region II covered the diagnostic fragments commonly used for the identification and structural elucidation of archaeal lipids. These product ions were derived from cleaving the ether bonds (C-O) of the glycerol tetraether structures. The ether bond cleavage resulted in the neutral loss of diphytanyl or phytanyl chains, along with the sequential loss of H_2_O, C_3_H_4_O, and C_3_H_6_O_2_ as observed in Region I. This process usually involves transferring one or two protons from the neutral loss molecules to the charged product ions^41^.

Region III (m/z < 300) contained ions from the fragmentation of isoprenoid chains and headgroups (Fig. S5). They are diagnostic for determining the structure of both components but often overlooked during manual identification. To mitigate low dot-product scores of scare diagnostic fragments during mass spectral matching, *in silico* reconstruction of Region III fragments was performed to improve annotation rates for targeted diether and tetraether lipids. These fragment ions are abundant and critical for improving the identification scores, which may be important particularly for lipids at low abundance that would otherwise evade detection due to insufficient spectral data.

After obtaining fragmentation patterns and relative ion intensities for a representative lipid, these data were used to reconstruct the MS^2^ spectra for its lipid class in a modeling effort that integrated known structural features of archaeal lipids, including characteristic headgroups, core structures, side-chain moieties, and cleavage patterns.

The above comprehensive workflow culminated in the ArchLips spectral library, which encompassed MS^2^ spectra for 199,248 archaeal lipids (Table 2), providing a robust resource for systematic identification and structural characterization of these molecules. To improve this library’s reliability, we compiled a high-confidence sub-dataset of 29,446 archaeal lipids by curating compounds reported in the literature, their derivatives, and lipids manually verified through rigorous laboratory identification. These entries were supplemented with *in silico* MS^2^ spectra to support robust spectral matching (Table 2). Additionally, we addressed challenges posed by automated peak-picking algorithms, which can misassign monoisotopic peaks due to interference from high molecular weight compounds or elevated [M+1] isotopologue signals (Fig. 3). To mitigate this challenge, we developed a sub-spectral library comprising 123,686 archaeal lipids, enhancing detection of compounds with [M+1]/M ratios exceeding 1, a threshold distinguishing true monoisotopic peaks from isotopic interference (Fig. S6). The spectral libraries were further strengthened by a prediction of the retention time based on the model built from manually curated dataset for our chromatographic condition (Fig. 1). The RMSE for the training dataset was 0.42 min, with a 95% confidence band of +/- 0.7 min, while the RMSE for the dataset was 1.29 min, and a 95% confidence band of +/- 2.11 min. These complementary strategies collectively refine lipid analysis accuracy by resolving peak assignment ambiguities inherent in automated workflows.

**Figure 3.**
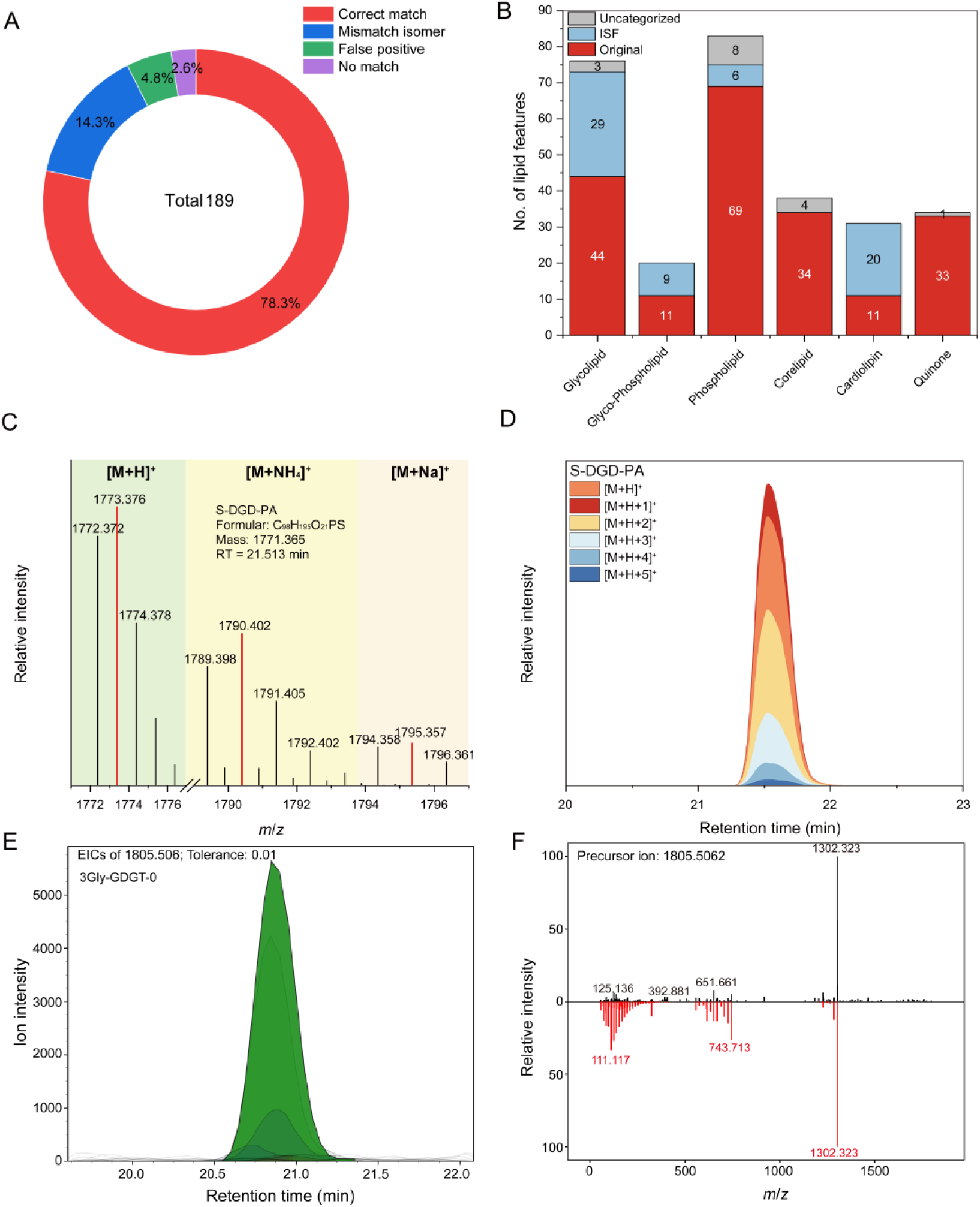
The performance of ArchLips in lipid identification of pure culture samples. A) The library search of 189 MS^2^ spectra of archaeal lipids from the peer-reviewed literature against the high confidence ArchLipid spectral library. B) The comparison of features of archaeal lipids in-source fragmentation (ISF) ions in quality control samples. C). Mass spectrum of a S-DGD-PA lipid showing the domination of M+1 isotopologues (red peak) arising from natural ^13^C occurred in some archaeal lipids. D) Distribution of chromatographic peaks of isotopologues in S-DGD-PA. E) Extracted ion chromatogram of 3Gly-GDGT-0 (*m*/*z* = 1805.5062, [M+NH_4_]^+^) in a M. *maripaludis* mutant strain and QC samples. F) Spectral matching of experimental MS² spectra of 3Gly-GDGT-0 (black, top) against referenced *in silico* spectra from the ArchLips database.

### Library validation using pure culture strains

To validate the accuracy of the ArchLips database, a decoy search of 189 MS^2^ spectra of archaeal lipids obtained from the peer-reviewed literature was performed against the high-confidence spectral library as described in Ref.^35, 59^. Over 78.0% of the MS^2^ spectra were correctly matched, 14.3% were annotated as homolog compounds and 7.4% as false positive or no match (Fig. 3A). The decoy searches of the high confidence spectral library and full spectral library against the non-archaeal-dominated public lipid spectral databases resulted in a relatively low hit rate for the GNPS (0.80%, 6.62%) and LipidBlast library (0.19%, 0.60%), respectively (Table S3).

Next, the performance of ArchLips database was evaluated using four pure cultures of archaea that have existing lipid profiles^24, 55, 60, 61^. The MS^2^ spectra matching results between the experimental spectra and the *in silico* spectral library are shown in Figures S7 and S8 (also including MS^2^ spectra detected in environmental samples). When applying the high-confidence spectral library without setting retention time (RT) for scoring, 236 mass features were successfully annotated with an identification score > 70% using MS-DIAL software. With an RT tolerance of 3 minutes, 198 mass features were annotated. Additionally, when using the high-confidence spectral library containing M+1 spectra, both with and without RT tolerance of 3 min, identified 260 and 231 mass features, respectively. These features were assigned to 112 to 131 unique archaeal lipid compounds, including diethers, tetraethers, cardiolipins, GDDs, and quinones (Fig. 4).

**Figure 4.**
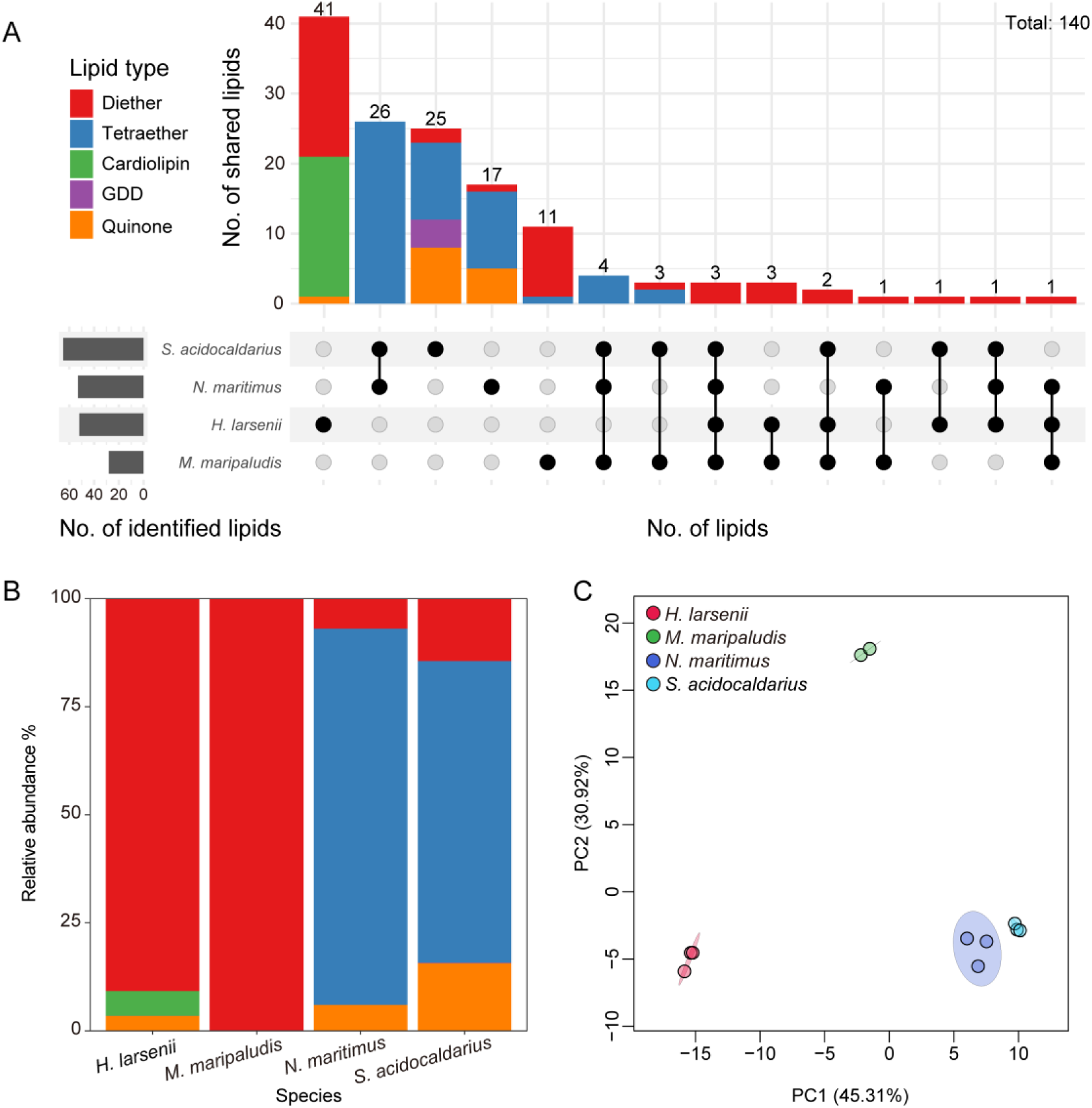
Application of the ArchLips database in lipid identification of archaeal strains. A) The intersection of lipid compounds of 140 identified from different archaeal species is illustrated using the UpSet plot. B) The lipid composition of major lipid groups in four archaeal strains. C) The PCA plot is based on the relative abundance of all identified lipids from the four pure cultures.

After manually verifying the lipid annotation results from the MS-DIAL software, an additional 9-28 unique compounds were identified, which together added to a total of 140 individual lipids across four archaeal pure culture strains (Fig. 4). Among them, only 3 diether lipids were shared by all four archaeal strains and 45 lipids were shared by subsets of these strains (Fig. 4A), which accounted for 32.1% of total lipid species. The remaining majority (67.9%) of the identified lipids were strain-specific. For example, the *N. maritimus* and *S. acidocaldarius* were both dominated by tetraether lipids, they were still distinguishable in the overall profile (Fig. 4C). Furthermore, *N. maritimus* contains methoxy archaeol (MeO-AR, *m*/*z* =667.6962, [M+H]^+^) that have been identified as an exclusive biomarker for Nitrososphaeria (Fig. 5A)^61^. The lipidome of *M. maripaludis* and *H. larsenii* were primarily composed of diether lipids, which were also distinct between these two strains. For *M. maripaludis* mutant strain, due to heterologous expression of Tes homologs, a trace amount of GDGT-0 (*m*/*z* = 1302.3197, [M+H]^+^)^22^ along with the intact polar form of 3Gly-GDGT-0 (*m*/*z* = 1805.5062, [M+NH_4_]^+^) were detected, albeit its wild-type strain did not produce GDGTs^22^ (Fig. 3).

A feature-based molecular network (FBMN) was constructed to characterize the spectral similarities between identified lipids and unknown features^62, 63^. The results revealed the FBMN contained a total of 770 nodes, which formed 97 clusters with at least two connected nodes (Fig. S9). These features in FBMN totally accounted for 28.9% of features with MS^2^ and 200 features among them were annotated using the ArchLips database. Figures 5A and 5B showed two clusters of archaeal diether and tetraether lipid-related features, respectively. The feature with a precursor ion at *m*/*z* 669.68 in Figure 5A was not annotated. As it had a mass difference of 324.12 Da from its adjacent node, 2Gly-Archaeol (20:0(OH)_20:0), which corresponds to two glycosyls, we manually assigned these two features as Archaeol (20:0(OH)_20:0). Besides, several chimeric mass spectra originating from the fragmentation of [M+1] isotopologue peaks were also observed, such as features at *m*/*z* 831.73 in Figure 5A and *m*/*z* 1457.3, *m*/*z* 1615.34 and *m*/*z* 1617.34 in Figure 5B. Additionally, the cluster of tetraether lipids in Figure 5B was primarily associated with calditol-GDGTs, which are suggested to be produced by thermoacidophilic archaea in the *Crenarchaeota* phylum (e.g., *S. acidocaldarius*) and the biosynthesis of these compounds requires a putative calditol synthase (cds)^60^.

**Figure 5.**
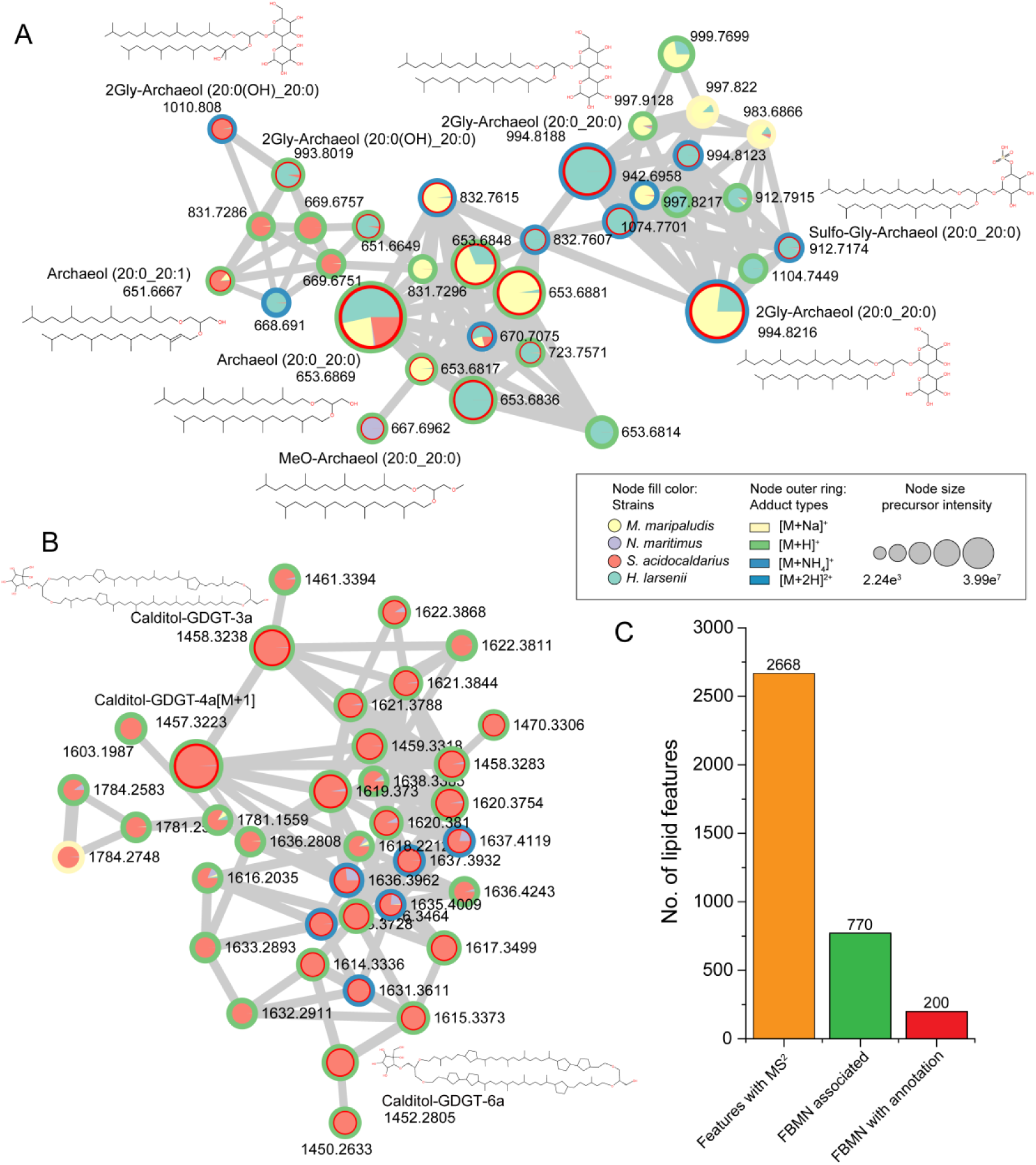
A subnetwork of MSMS spectral similarity between annotated features with ArchLips database and unknown features in archaeal pure cultures. A) diether lipids; B) tetraether lipids lipid features involved in the FBMN. The filled color of the nodes indicated the distribution of lipid features in four archaeal strains. The color of the inner rings represents annotated features (red), and the color of the outer rings refers to the adduct types. The width of the line connecting the nodes represents the level of mass spectral similarities, with thicker lines indicating greater similarity between two nodes. C) the bar chart showing the features with MS^2^ and associated in FBMN and annotated features retained in FBMN.

### Archaeal lipidome applied to diverse natural habitats

To identify archaeal lipids in environmental samples using the ArchLips database, 52 samples were collected from diverse geological settings and habitat types such as surface and subsurface marine sediments, cold seep sediments, hot spring sediments, acid mine drainage (AMD), aerated soils and permafrost soils. A range of 415 to 551 archaeal ether lipid features (assigned to 176 to 249 lipid compounds) and 47 to 56 non-ether lipid features (assigned to 20 to 34 lipid compounds) were detected using the untargeted spectral libraries (Fig. S10), which was 1-2 orders of magnitude greater than results obtained from targeted approaches in any environmental samples. Comparison among these samples showed first-order differences in lipid compositions between marine and terrestrial environments (marine-terrestrial “lipid divide”) (Fig. 6A).

**Figure 6.**
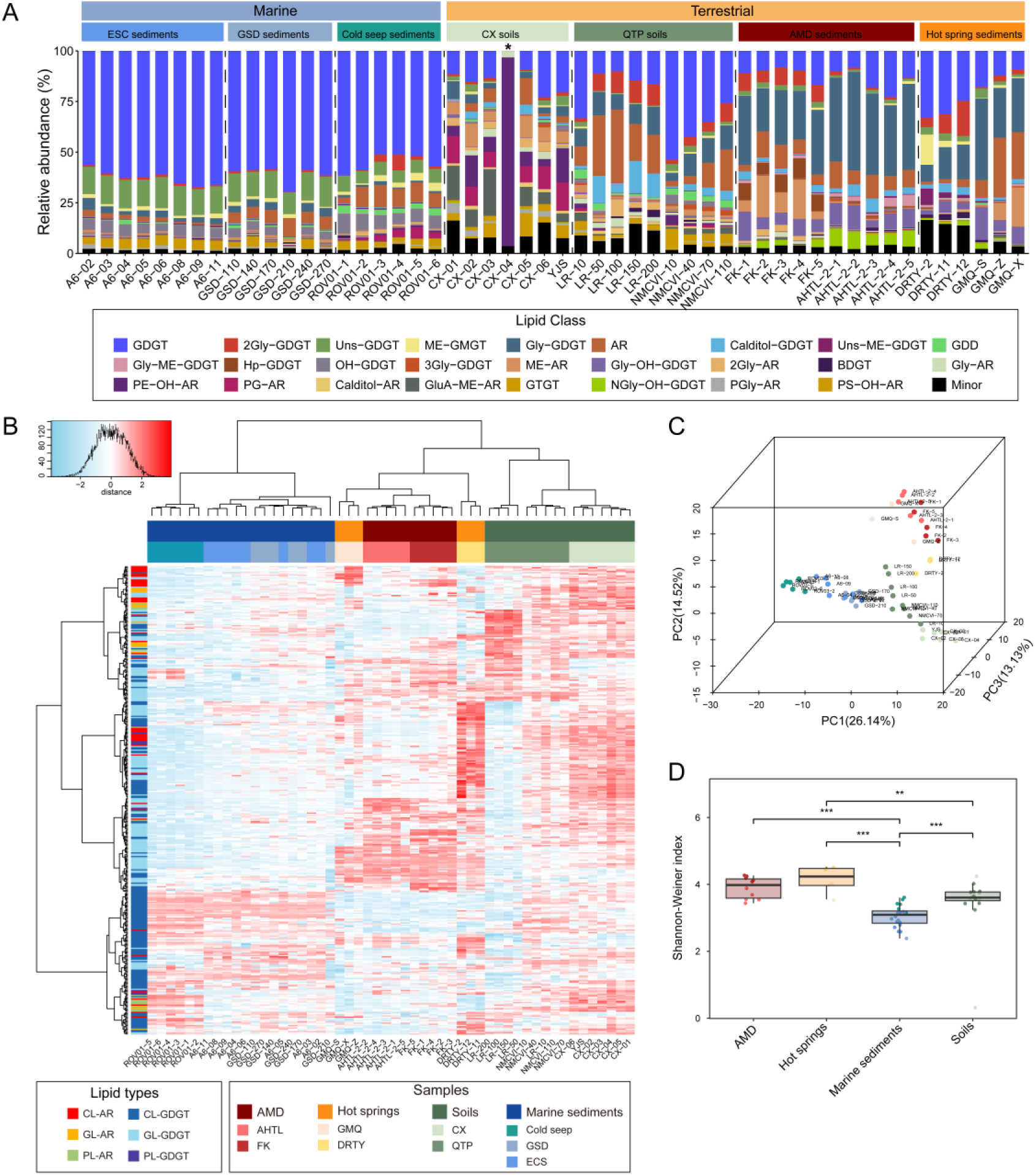
Lipidomic profiles of archaeal lipids from environmental samples identified using ArchLips database. A)Lipid composition, B) Clustering analysis using Euclidean distance and Ward.D method, C) PCA analysis and D) Shannon diversity of archaeal lipids. AMD, acid mine drainage. AHTL and FK represent samples collected from Tongling and Fankou AMD, respectively. GMQ and DRTY represent sediment samples collected from hot springs in Tengchong. ROV01 represents cold seep sediment collected from the South China Sea, GSD a sediment core from Pearl River estuaries and ESC surface sediment samples from the East China Sea. CX and QTP represent soil samples from the western region of Sichuan Province and Qinghai-Tibetan plateau permafrost, respectively.

Clustering analysis based on the relative abundance of archaeal lipids further depicted the habitat types influencing the marine-terrestrial “lipid divide” (Fig. 6B and Fig. S12). Among marine samples, moderate seepage (ROV01) samples formed a separate branch from surface and subsurface sediments of typical marine environments. In the terrestrial group, samples from extreme environments, such as hot spring sediments and acid mine drainage (AMD), clustered together. Notably, lipid composition varied significantly between acidic and alkaline hot spring samples, with alkaline hot springs clustering with AMD samples. This pattern aligns with differences observed in 16S rRNA-based archaeal community composition between acidic and alkaline hot springs (Fig. S13). Soil samples formed separate branches, further dividing into subgroups of aerated soils (CX and NMCVI) and wet permafrost soils (LR), mirroring soil water content or oxygen availability driven archaeal adaptations^64^.

The archaeal lipid composition was also evaluated using principal component analysis (PCA) (Fig. 6C). The first three principal components explained 53.79% of the variance in archaeal lipids, with samples from different biomes primarily separating, consistent with the hierarchical clustering results (Fig. 6B). We further assessed the Shannon-Weiner diversity of archaeal lipidomes (Fig. 6D). Hot spring samples exhibited the highest lipid diversity, followed by AMD samples. At the same time, soil and marine sediment had the lowest diversity. However, this pattern does not fully align with archaeal community diversity inferred from 16S rRNA gene analysis, where AMD samples and marine sediments showed the highest diversity and soils had the lowest (Fig. S14).

## Discussion

We present the first comprehensive database ArchLips that is constructed using a combinatorial and heuristic algorithm, which should fill a critical gap in existing lipidomic resources. It was meticulously developed to enhance archaeal lipid identification in four key aspects: (1) comprehensiveness, (2) accuracy, (3) efficiency, and (4) enhanced in biological interpretation. These components are detailed below:

1. *Comprehensiveness*. Although previous studies have reported diverse structures of archaeal lipids, this information remains fragmented within the literature^31,65^. Our database curated and integrated these data into a comprehensive repository that now constitutes 218,868 archaeal lipid structures, which includes a complete range of core lipids and a wide range of intact polar lipids (IPLs) (Fig. 1 and Fig. S1). These structures allow us to generate over 380,000 *in silico* MS^2^ spectra of two adducts for each lipid to improve lipid identification. For verification, among 189 MS^2^ spectra reported in the literature, 92.2% of them matched those *in silico* MS^2^ spectra in our libraries. This new resource should enable precise lipid searches using accurate-mass-based methods common in shotgun lipidomics, support structure-driven molecular-level lipidomics through untargeted MS^2^ spectral matching paired with in silico databases, and allow direct linking of molecular identities to their chemical structures.
2. *Accuracy*. ArchLips demonstrated high accuracy in identifying archaeal lipids by matching 189 curated MS² spectra from peer-reviewed literature against the high-confidence spectral library (Fig. 3). Decoy searches further validated the database’s reliability, while low hit rates in LipidBlast and GNPS comparisons (Table S3) underscored its specificity for archaeal lipid structures. Validation using pure archaeal cultures and environmental samples confirmed its applicability across diverse settings (Figs. S6 and S7). Additionally, ArchLips enhances identification confidence through reproducible, efficient spectral libraries, minimizing reliance on manual interpretation, which is often influenced by analyst expertise, thereby improving consistency and accuracy.
3. *Efficiency*. Compared to targeted approaches, untargeted mass spectrometry-based lipidomics significantly reduces the time required for sample preparation while enabling comprehensive detection of a broad range of lipid species^58, 66^. The efficiency of our database is characterized by being capable of batch identification of hundreds of archaeal lipid species from a sample (Fig. 4 and Fig. S10) through cheminformatic tools such as MZmine^67^ and MS-DIAL software^68, 69^. This allows us to streamline data processing pipelines (Fig. S2) and to calculate lipid-based chemotaxonomic, paleoecological, and paleoclimate proxies (Fig. S15). Furthermore, the ArchLips can be integrated with other public lipidomic databases (e.g., LipidBlast), supporting comprehensive, cross-domain characterization of archaeal, bacterial, and eukaryotic lipids (Fig. S11).
4. *Enhanced in biological information*. Increasing studies have demonstrated that the clustering tree based on archaeal lipids closely resembles the topology of the 16S rRNA gene tree^24, 70, 71^. This study integrated global lipid annotation with feature-based molecular networking (FBMN) to reveal strain-level structural diversity in archaeal lipids (Fig. 5C), which supports previous findings (Elling et al., 2017; Yao et al., 2023). We also applied the ArchLips full spectral library for high throughput lipid identification to a lipidomic dataset of four halobacterial strains analyzed on a Q Exactive Orbitrap MS (ftp://massive-ftp.ucsd.edu/v04/MSV000089830/)^72, 73^. The raw data were processed using MS-DIAL software, leading to the successful annotation of 219 lipid features, which enabled a direct comparison of lipid compositions between different strains. These lipids represented a broad spectrum of archaeal lipid classes that included both previously reported^72, 73^ and novel species (Fig. S16), indicating the ArchLips database can be used for lipid identification across different mass spectrometry platforms.

The analysis of 52 environmental samples indicated that archaeal lipid composition reflects differences in habitat type, such as marine vs. terrestrial, extreme vs. mesophilic environments, which to some extent align with the archaeal community structures based on 16S rRNA sequencing (Fig. S12). However, direct linking archaeal lipidomes to 16S rRNA-based community composition remains challenging and requires careful evaluation of confounding factors such as DNA and lipid degradation, as well as differentiation of sources (e.g., terrestrial input vs. water-column-derived lipids deposited in sediments).

While ArchLips represents a significant advancement for high-throughput identification of archaeal lipids, several key limitations warrant should be careful under consideration. The caution for using automated lipid identification lies in the presence of structural isomers and in-source fragmentation, which may cause false identification. The properties of archaeal lipids, especially tetraethers, such as GDGTs, can form a variety of structural isomers due to the flexible positional combinations of a number of cyclopentane (and sometimes cyclohexane) rings and double bonds along the C_40_ isoprenoid chains^43, 44, 47, 49, 52^. For example, GDGTs containing cyclopentane rings can be mistakenly assigned as unsaturated GDGTs, as they are fragmented into identical product ions that may only vary in ion intensity and retention time^52^. These isomers cannot be resolved using tandem mass spectra or reverse phase chromatography, but some of them may be resolved under rigorous normal phase chromatographic conditions and often assisted by manual verification^43, 44, 55^.

Another caveat is the inherent nature of the electrospray ionization (ESI) technique that causes in-source fragmentation (ISF) of some molecules during the initial ionization process and transports the fragmented ions into the collision cell^74^. Despite extensive efforts to minimize ISF through the optimization of mass spectrometry parameters, it remains a challenge and can lead to false identifications of target compounds. In this study, 80 annotated MS features were likely to be ISF from the pure archaeal cultures, representing 33.8% of the total annotated features. This large proportion of ISF can impact the relative abundance of a compound susceptible to be the product of a fragmenting molecule; however, they do not appear to affect the identification of true archaeal lipids using our quadrupole time-of-flight high-resolution mass spectrometry (qTOF-HRMS) platform. Further analysis showed that ISF predominantly occurred in glycolipids, glyco-phospholipids, and cardiolipins, suggesting that greater caution needs to be exercised when estimating the abundance of these lipid classes. One effort to mitigate the false identifications caused by ISF and structural isomers is to incorporate retention time (RT) as an additional discriminative parameter. Since RT is highly sensitive to experimental conditions, it requires consistency in the chromatograph conditions used to generate spectral libraries, which may be improved by developing RT prediction models based on machine learning or artificial intelligence (AI) (Fig. 3).

Overall, our archaeal lipidomic results provide a more comprehensive reflection of archaeal community diversity than previous methods and demonstrate promise for establishing high-resolution lipid-based ecological indicators and paleoclimate proxies, which may help open up new research directions such as lipo-stratigraphy^75, 76^ or paleo microbial ecology into deep geological time when molecular DNA becomes less accessible or non-existing.

## Methods

### Archaeal cultures and sediments

Two wildtype pure cultures and two mutant strains of archaea were examined in this study, which included a *Nitrosopumilus maritimus* strain SCM1^77^, a *Haloferax larsenii* JCM 13917, a *Methanococcus maripaludis* mutant strain pMEV4-ma_1486 and a mutant strain of *Sulfolobus acidocaldarius* (p1561-grsB-grsA). *N. maritimus* is the first pure strain of marine group I Thaumarchaeota (now the class Nitrososphaeria)^77^, which grows chemolithoautotrophically and obtains energy by performing aerobic ammonia oxidization, thus called ammonia-oxidizing archaea (AOA). The lipidome of this strain has been extensively examined and covers a suit of archaeal tetraether lipids, including the characteristic biomarkers crenarchaeol and its regio-isomer crenarchaeol for AOA^30, 54, 78^. *H. larsenii* and *M. maripaludis* are two euryarchaeotes belonging to the class Halobacteria and methanogens, respectively. The strains of these two euryarchaeotes do not produce tetraether lipids, but diethers as the major lipid content^24^. In this study, a *M. maripaludis* mutant was used, which produces GDGT-0 through the expression of *Tes* homologs^22^. *S. acidocaldarius* is a thermoacidophile belonging to Crenarchaeota, which is capable of producing a high diversity of tetraether lipids, including the Calditol-GDGTs and GDGTs up to 8 cyclopentane rings. The *Sulfolobus acidocaldarius* mutant strain (p1561-grsB-grsA) was used for examining enhanced biosynthesis of GDGTs with more cyclopentyl rings (e.g., GDGT-5 to GDGT-8).

The four archaeal strains were cultivated according to previously published cultivation protocols. Briefly, the *N. maritimus* strain SCM1 was aerobically cultured at 30 ℃ with 1L Synthetic Crenarchaeota Medium (SCM) as described in Ref. ^77^. The cells were harvested at the stationary phase by filtering through a 0.22 μm PVDF membrane filter. The *M. maripaludis* S001 mutant (pMEV4-ma_1486) strain was grown in McF medium under a gas stream of N_2_/CO_2_ (80:20) at 37 °C as described in Zeng et al. ^22^. The *S. acidocaldarius* mutant (p1561-grsB-grsA) strain was cultured in Brock medium following Zeng et al. ^55^. *H. larsenii* (JCM 13917) was grown in triplicate in DSMZ 589 liquid medium at 37 °C and pH 7.4 with 200 rpm shaking, as described in Yao et al. ^24^. Biomass of the latter three strains at the stationary phase was collected by centrifugation at 10000 g for 10 min.

This study also examined a total of 52 sediment and soil samples across a wide range of environmental gradients, including sediment from hot springs in Tengchong, Yunan Province, sediment from acid mine drainages in Anhui and Guangdong provinces^79^, permafrost soil from Tibet Plateau^80^, soil from Western Sichuan Plateau, surface sediment of cold seeps^81^ and a sediment core from the South China Sea^82^, and sediment from the East China Sea^83^. These samples were collected from 2015 to 2023 and stored at −20 ℃ in the laboratory until analyses. Detailed sample information is shown in Table S1.

### Lipid analysis

Total lipid extracts (TLEs) of all samples were prepared using the Bligh & Dyer method modified by Sturt et al. ^84^. Briefly, freezing dried biomass or environmental samples were extracted twice using a combination of methanol (MeOH): dichloromethane (DCM): phosphate buffer (PB) (2:1:0.8, v/v/v) and subsequently extracted twice with a solvent mixture of MeOH: DCM: trichloroacetic acid buffer (TCA) (2:1:0.8, v/v/v). During each extraction, samples were sonicated for 10 min and then centrifuged at 3000 rpm for 5 min. The supernatants were combined, and the solvent ratio was adjusted to 1:1:0.9 (v/v/v) by adding an extra amount of DCM and ddH_2_O. After phase separation was achieved, the DCM layer was collected and the remaining aqueous was re-extracted twice with DCM. Finally, the DCM phase was combined, dried under a gentle nitrogen gas stream, and stored at −80 ℃ until further analysis.

An aliquot of the TLEs was dissolved in methanol for injection. Archaeal lipids were separated using a ACQUITY I-Class Ultra performance liquid chromatography (UPLC) on a C18 EXCEL UPLC column (2.1×150 mm, 2 μm; ACE) maintained at 45 ℃ with a reverse phase gradient. The eluting gradient was modified according to Zhu et al. ^66^ : 0-5 min 100% A, 5-10 min 0-24% B, 10-36 min 24-60% B, 36-38 min 60-90% B, 38-45 min 90% B, 45-45.1 min 90-100% A, 45.1-55 min 100% A. Solvent A (Methanol) and solvent B (isopropanol) are both amended with 0.1% NH_4_OH (25–30% NH_3_ basis) and 0.04% formic acid (>99.0%). The flow rate was 0.3 ml/min and the total run time was 55 min.

MS data were acquired in a Resolution mode by a Waters SYNAPT G2-S*i* quadrupole time-of-flight mass spectrometer (qTOF) coupled to an electrospray ionization (ESI) source operated at positive ion mode. The MS source parameters were identical to Chen et al. ^85^: capillary 2.5KV, source temperature 120 ℃, source offset 80 V, sampling cone 45V, desolvation gas 800 L/hr at 350 ℃, cone gas 50 L/hr, nebulizer 6.5 bar. The mass range for MS^1^ and MS^2^ was set to 100 - 2000 and 50 - 2000, respectively. The MS^2^ spectrum was acquired by FAST-DDA mode, and the top 5 most abundant ions were selected for MS^2^ via collision-induced dissociation (CID) with a mass-dependent ramped collision energy that started at 10 V, and ended at 15 V for low mass (*m*/*z* 100); and started at 55 V, ended at 65 V for high mass (*m*/*z* 2000). The qTOF mass spectrometer was calibrated with sodium iodide (*m*/*z* 50–2000; Residual mass error <0.5 ppm) and a real-time calibration with leucine enkephalin solution ([M+H]^+^, m/z 556.2771).

### Data processing

The raw data obtained by qTOF were converted into mzML format with the Waters2mzML script (V1.2.0)^86^ and the mzxml-precursor-corrector script was employed to fix the precursor values^87^. The processed mzML files were imported into the MS-DIAL software for data processing. Parameters included a minimum peak height of 1000 amplitude and mass slice width of 0.1 Da. The peaks were aligned with a retention time tolerance of 0.3 min and an MS^1^ tolerance of 0.05 Da. Lipids were annotated with mass error within 0.01 Da for MS^1^ and 0.05 Da for MS^2^, and total scores over 70% that matched the library were considered as identified compounds. A cutoff at 1% MS^2^ spectrum relative abundance was used for compound identification to reduce the effect of false positives caused by [M+Na]^+^ ions. The feature table exported from the MS-DIAL software can be further processed using our python script to find possible adducts of identified features and, subsequently, the peak area, which can be combined into a lipid compound for proxy calculation.

### Feature-Based Molecular Networking Construction

A molecular network was created with the Feature-Based Molecular Networking (FBMN) workflow on GNPS^63, 88, 89^. The mass spectrometry data processed with MS-DIAL were exported to GNPS for FBMN analysis. The data was filtered by removing all MS^2^ fragment ions within +/- 17 Da of the precursor *m*/*z*. The MS^2^ spectra were window-filtered by choosing only the top 6 fragment ions in the +/- 50 Da window throughout the spectrum. The minimum fragment ion intensity in the MS^2^ spectra was set to 100. The precursor ion mass tolerance was set to 0.01 Da, and the MS/MS fragment ion tolerance to 0.05 Da. A molecular network was created where edges were filtered to have a cosine score above 0.7 and more than 6 matched peaks. Further, edges between two nodes were kept in the network if and only if each of the nodes appeared in each other’s respective top 10 most similar nodes. The maximum size of a molecular family was set to 100, and the lowest-scoring edges were removed from molecular families until the molecular family size was below this threshold. The molecular networks were visualized using Cytoscape software. Network visualization was performed using Cytoscape software (https://cytoscape.org/).

### Curation of the ArchLips database

#### *In silico* molecule structures

The core structures of tetraether lipids were manually drawn using the MarvinSketch software (V23.16.0), while the core structures of diether lipids were generated using the ChemAxon Reactor software (V21.3.0). The core structures were stored in EXCEL files to be extended or modified with JChem for EXCEL software. Additional structural information, including molecular formula, exact mass, SMILES, Inchi, InchiKey, and logP were also calculated using the RDKit package (https://www.rdkit.org) in Python (https://www.python.org/).

The structures of intact polar lipids were generated by connecting the core structure templates with diverse polar headgroups. A total of 79 head groups were obtained from the literature, (e.g., Ref. ^23, 41–54^, LipidMAPS (https://www.lipidmaps.org/) and LipidBank (https://lipidbank.jp/) were taken under consideration, including 39 phosphoric groups, 38 glycosyl groups (including a calditol group), one methylated/ acetylated group were also included (Fig. S1 and Table S2). Their structure (MDL molfiles) and structural information were also created and stored in an EXCEL file. The structures of intact polar lipids (IPLs) were created based on combining the SMILES with the RDKit package in Python. The generated SMILES could then be transformed into the MDL molfile structures, with the structural information (e.g., formula, exact mass, InChI, InChIKey, logP) calculated using the RDKit package and stored in SDF files. The MDL molfile structural files of IPLs were further standardized by the RDKit package using core lipids as a template. The structure of cardiolipins and quinones was constructed following a similar protocol (Fig. 1).

#### *In silico* MS^2^ spectral Library

A modified LipidBlast template was used for generating the *in silico* spectra library^90^, incorporating two main precursor ions with adducts of [M+H]^+^ and [M+NH_4_]^+^, commonly detected under positive ion mode. This template preserved all structural information on a dedicated sheet, with MS^2^ data organized into multiple EXCEL sheets, each corresponding to a lipid class with similar core lipids. For example, all the PG-archaeol homologs with different unsaturation and chain lengths were stored in the same sheet. The first 28 columns of each MS^2^ sheet included the molecule name, structural information, and sidechain properties, followed by columns recording the *m*/*z* and intensity of MS^2^ peaks. The core lipid MS^2^ sheet served as a template, recording original sidechain properties shared by subsequent sheets, with other MS^2^ sheets synchronizing data from this template.

Structural information for each MS^2^ sheet was extracted from the structure sheet using the built-in VLOOKUP function of EXCEL, allowing updates and expansions by editing the structure sheet templates. The MS^2^ data and structural information were then exported as MSP-formatted library files, divided into 17 files according to lipid type to reduce the computational burden. The library exportation program, rewritten in python, employed multiprocessing to handle multiple files simultaneously, expediting the process. During exportation, peak intensities for each molecule were standardized by normalizing the highest peak intensity to 1000 and removing peaks with intensities lower than 0.1. The exportation process for approximately 400,000 MS^2^ spectra took less than 8 minutes, resulting in an MSP file of 1.46 GB.

#### Parameters for library validation

A total of 189 tandem mass spectra (MS^2^) of archaeal lipids and the characteristic peaks in MS^2^ reported in previous publications were digitalized using the WebPlotDigitizer (v4.6)^91^. Those spectra were then formatted into an MGF file and annotated with our full lipid library and high-confidence library. The LipiDex (v1.1) was used for library validation^59^. All MS^2^ data to be tested in this part (our archaeal libraries, open sources libraries of LipidBlast and GNPS, and 189 archaeal lipid MS^2^ from literature) were transformed into brief versions of MSP files or MGF formatted files. The search parameters were set as ± 0.01 Da of MS^1^ or MS^2^ tolerance. The peaks with *m*/*z* > 61 in MS^2^ spectra were retained for similarity scoring.

#### The retention time (RT) prediction

The RT prediction was performed by Retip package (v0.5.4)^92^ in R (v4.3.1, https://www.r-project.org/). A dataset contained 265 identified lipids from the environment and pure culture samples were randomly divided into a training set and testing set (4:1). Five different algorithms (Random Forest, BRNN, XGBoost, lightGBM and Keras) were used to build prediction models based on the training set, and their prediction performances were calculated and compared with the testing set. The model with the lowest RMSE (root-mean-square error) value (the BRNN model) was used to predict RT for the *in silico* database.

#### Decoy search

The MSP formatted LipidBlast library (positive, version 68) was downloaded from http://prime.psc.riken.jp/compms/msdial/main.html#MSP, which included 81 classes, 377,313 molecules, and 554,041 spectra. The full GNPS library was downloaded from https://ccms-ucsd.github.io/GNPSDocumentation/gnpslibraries/, which covered 591,512 molecules. Both libraries do not contain archaeal lipids, so the annotation can confidently be categorized into a false positive result. Further, the libraries were annotated against themselves to evaluate the replicate spectra.

## Data availability

The converted raw data (mzML format) of archaeal lipidome of pure cultures and environmental samples used in this study is available in MassIVE under the accession code MSV000097125 and MSV000097126.

## Code availability

All custom scripts to generate the ArchLips spectral library in this study are available at https://github.com/magicthree/archaeal_lipid_MS2_library.

## Acknowledgments

We would like to thank Wenxiu Wang for providing GSD samples, Xiaotong Tang for the QTP permafrost soils and Bu Xu, Dongyu Cui for helping to collect soil samples from the western region of Sichuan Province. This research was supported by the National Natural Science Foundation of China (Nos. 32393974, 42372354 and 42003063), the Stable Support Plan Program of Shenzhen Natural Science Fund (20200925173954005), the Guangdong Basic and Applied Basic Research Foundation (2021B1515120080); the Shenzhen Key Laboratory of Marine Archaea Geo-Omics, Southern University of Science and Technology (ZDSYS201802081843490), and the High-Level Special Fund of SUSTech (G03050K001). Computation in this study was supported by the Centre for Computational Science and Engineering at the Southern University of Science and Technology. This paper contributes to the Science Plan of the UN Ocean Decade Global Ocean Negative Carbon Emissions (Global ONCE) Program.

## Author Contributions

F.Z. and C.Z. conceptualized the project, designed and oversaw the experiments, and wrote and revised the manuscript. W.Y. and W.H. performed coding and validation of the scripts, created the database, grew cell cultures, analyzed mass spectral data, and contributed to the writing and revision of the manuscript. W.Z., Y.C., and H.C. performed lipid extraction and lipid structure generation and contributed to the writing and revision of the manuscript. L.H. and Y.Z. helped with environmental sample collection and participated in the manuscript writing. Z.Z., X.L., Y.Z. and S.D. provided comments and advice and helped revision of the manuscript. All authors have reviewed and approved the submission of this manuscript.

