## Supplemental information for "ArchLips: A comprehensive *in silico* database for high-throughput identification of archaeal lipids"

**Supplementary Text**
**Supplementary Figures 1-16**
**Supplementary Tables 1-3**

### Supplementary Text

#### DNA extraction and sequencing

Total DNA was extracted from about 0.5 g of sediment or soil using FastDNA SPIN for soil kit (MP Biomedicals, LLC). The V4 region of prokaryotic 16S rDNA was amplified by PCR with universal prokaryotic primers (515F: GTGYCAGCMGCCGCGGTAA and 806R: GGACTACNVGGGTWTCTAAT) [1]. The 50 μl PCR mixture contained 25 μl of 2 ×Premix Taq DNA polymerase (TaKaRa), 0.2 mM of each primer, 3 μl of template DNA and $ddH_2O$. Thermocycling process included the following steps: initial denaturation for 30 s at 94 °C, followed by 30 amplification cycles consist of 30 s denaturation at 94 °C, 30 s annealing at 58 °C and 30 s elongation at 72 °C. The PCR products were pooled and purified by EZNA Gel Extraction Kit (Omega, USA) and sent to Guangdong Magigene Biotechnology Co., Ltd. for sequencing (Illumina Miseq).

The raw sequencing data of most environmental samples have been analyzed and published, including acid mine drainages in Anhui and Guangdong provinces[2], permafrost soil from Tibet Plateau[3], surface sediment of cold seeps[4] and a sediment core from the South China Sea[5], and sediment from the East China Sea[6].The raw sequencing reads were reanalyzed using the Quantitative Insights into Microbial Ecology (QIIME2, version 2023.5) software with default setting [7]. Taxonomy was assigned using the SILVA v138 99% dereplicated reference database (https://www.arb-silva.de/) [8].

**Supplementary Figures 1-16**

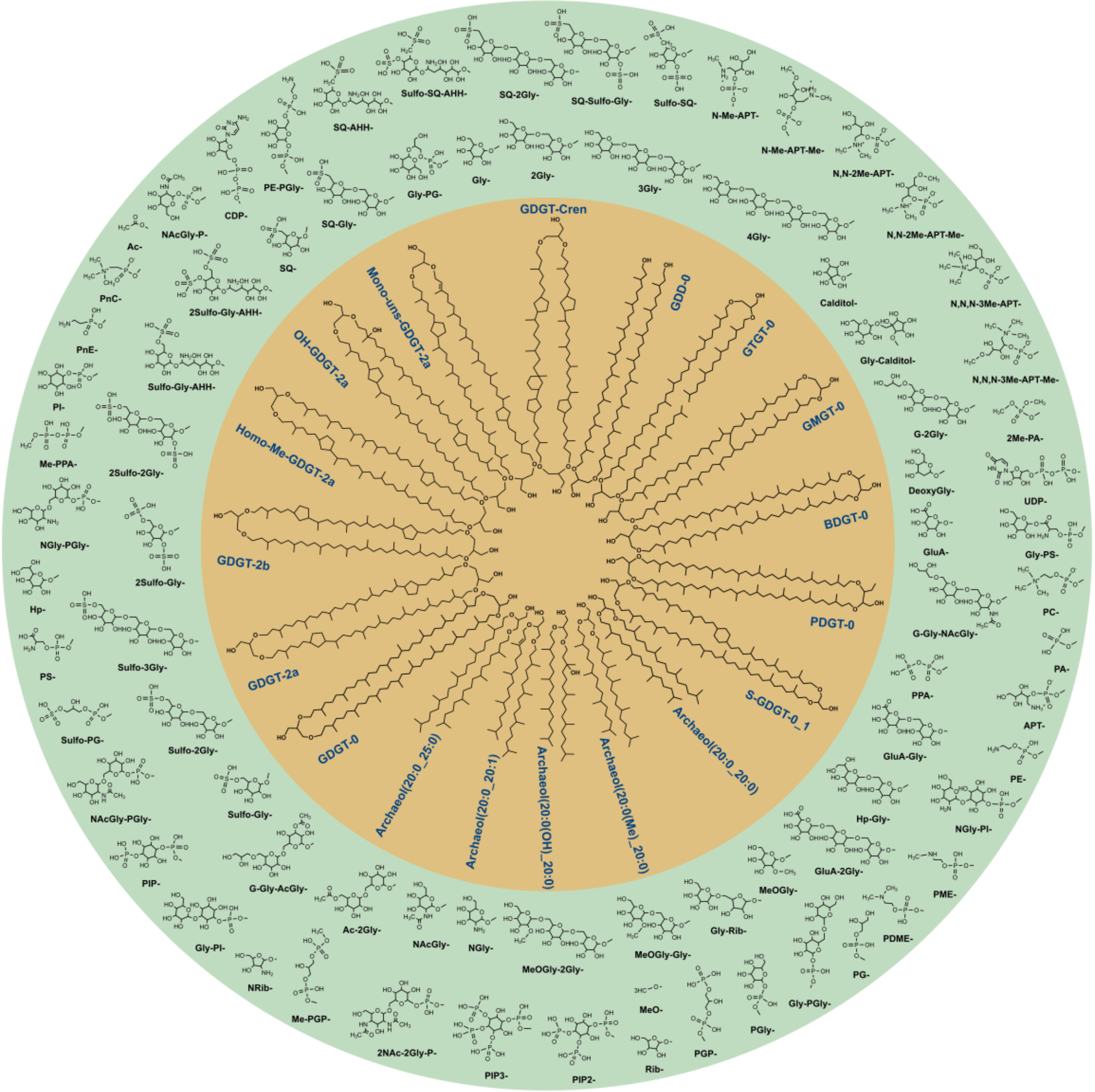

Figure S1. Representative molecules of archaeal lipids contained in the ArchLips database. GDGT refers to glycerol dialkyl glycerol tetraether. GDGT-cren, Crenarchaeol; BPG, bisphosphatidyl glycerol; S-GDGT, six-membered ring GDGT; PDGT, pentanetriol GDGT; BDGT, butanetriol GDGT; GDD, isoprenoidal glycerol dialkanol diether; GMGT, glycerol monoalkyl glycerol tetraether; GTGT, glycerol trialkyl glycerol tetraether. The full name of the polar head group is listed in Table S3.

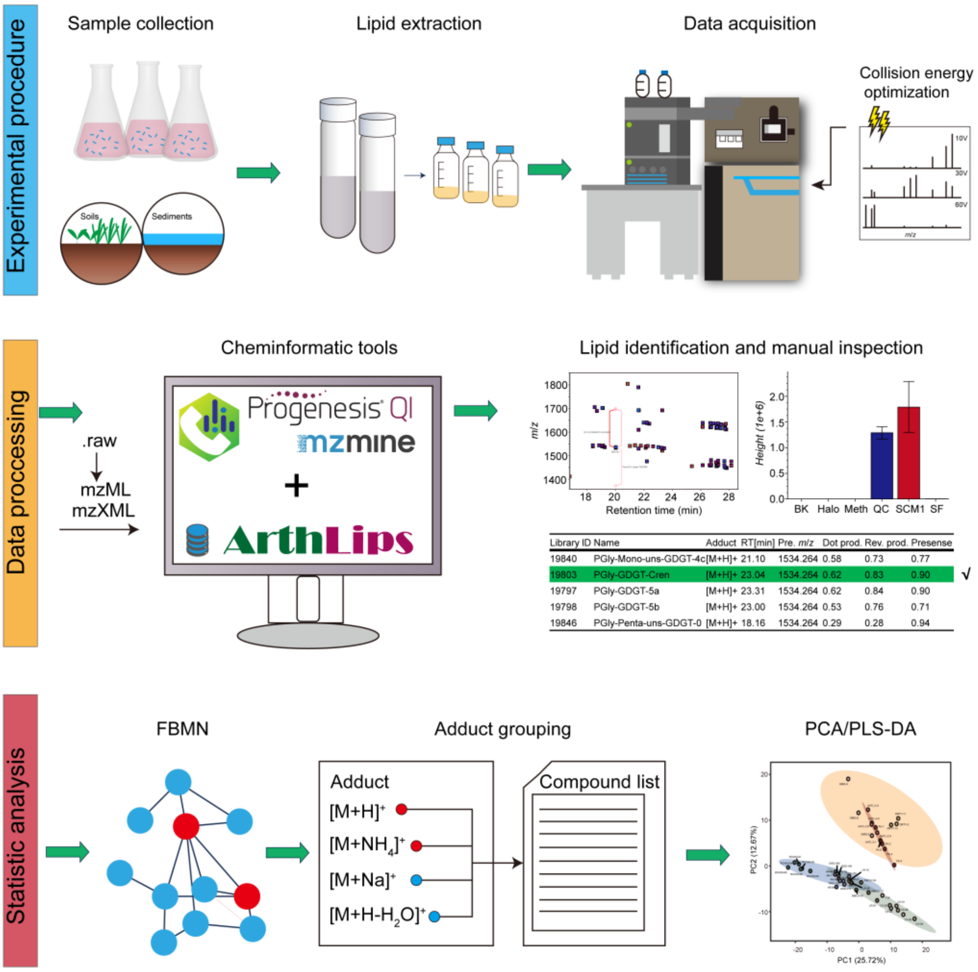


Figure S2. Workflow for archaeal lipid analyses using untargeted high resolution mass spectrometer (HRMS) couped to bioinformatic tools and the ArchLips database. Samples were collected from either natural environments or lab pure culture experiments. The lipid extraction was performed using a modified Bligh & Dyer method. The obtained total lipid extracts (TLEs) were analyzed on a reverse phase ultra-performance liquid chromatography quadrupole time of flight mass spectrometer (RP-UPLC-qTOF-MS). A MS/MS experiment was performed to obtain the optimization collision energy before data acquisition. The data processing was performed with commercial or accessed cheminformatic tools, and lipid identification was achieved using the ArchLips database and further manual inspection, followed by the building of feature-based molecular networking (FBMN), adduct grouping, and statistical analysis.

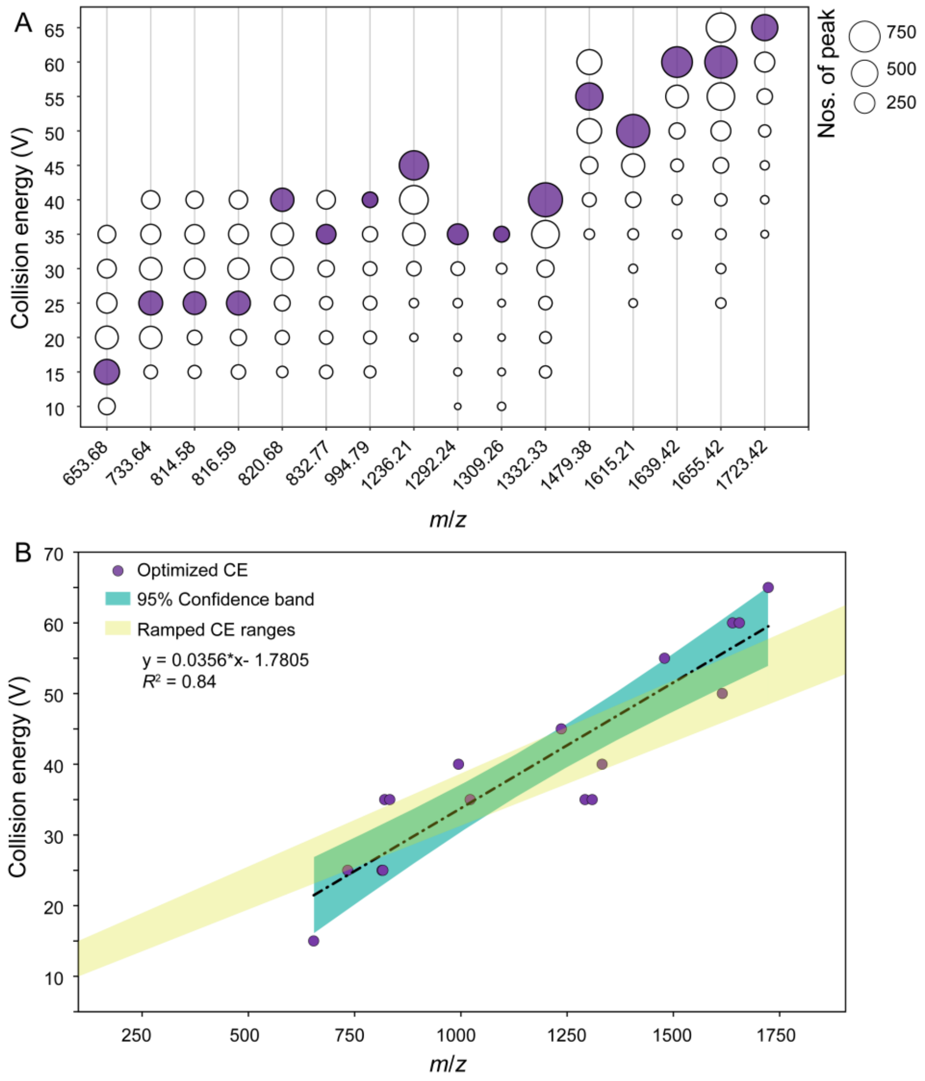


Figure S3. The collision energy (CE) optimization for collision-induced disassociation (CID) of archaea lipids. A) The collision energy range was used for cleavage of archaeal lipids; the color (red) of the circles represents the optimized CE values and sizes of the circles indicate the number of product ions with abundance > 0.001% of total peaks in $MS^2$ spectra; B) Regression of optimized CE verse *m*/*z* of precursor ions. The optimal energy was chosen as the one that produces the highest number of mass spectral peaks at different collision energies. A ramped CE range was used for data acquisition during the library construction.

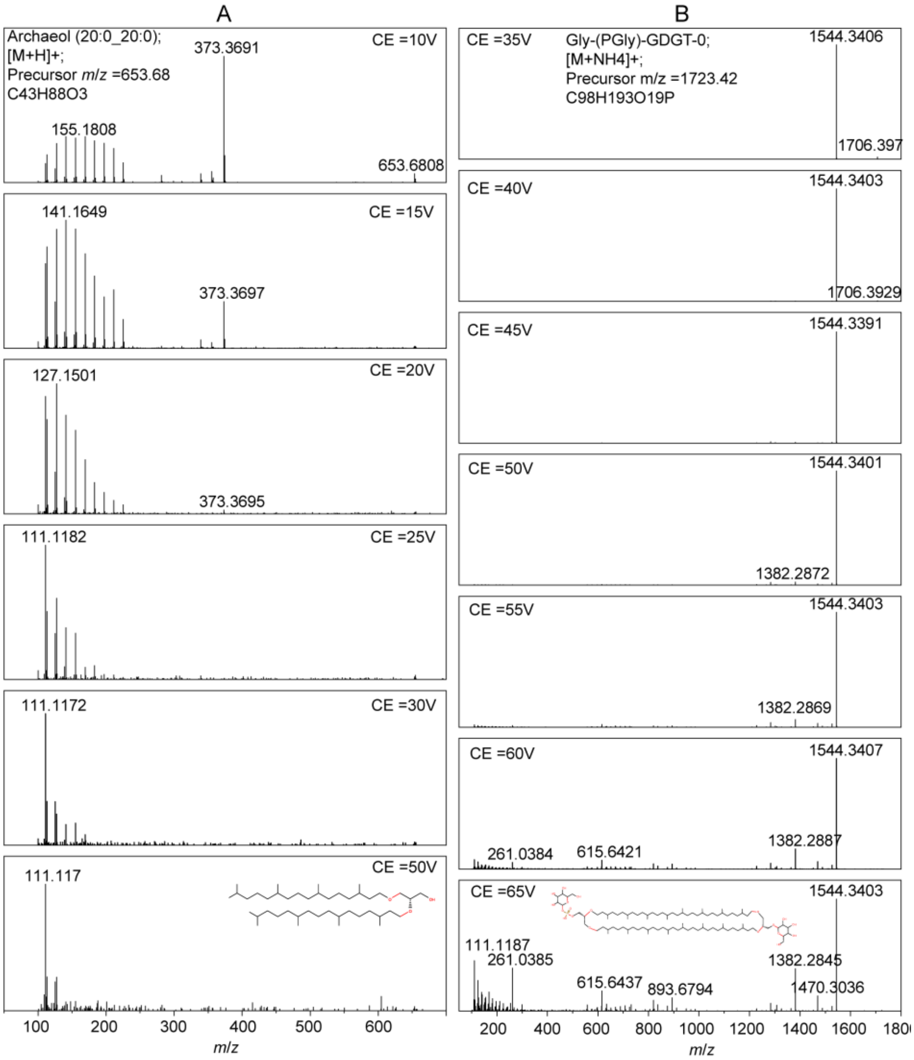


Figure S4. MS$^2$ spectra of archaeal lipids obtained under different collision energies (CEs) during CID experiments. A) Archaeol (20:0_20:0), $[M+H]^+$ at *m/z* 653.68; B) Gly-(PGly)-GDGT-0, $[M+NH_4]^+$ at *m/z* 1723.42.

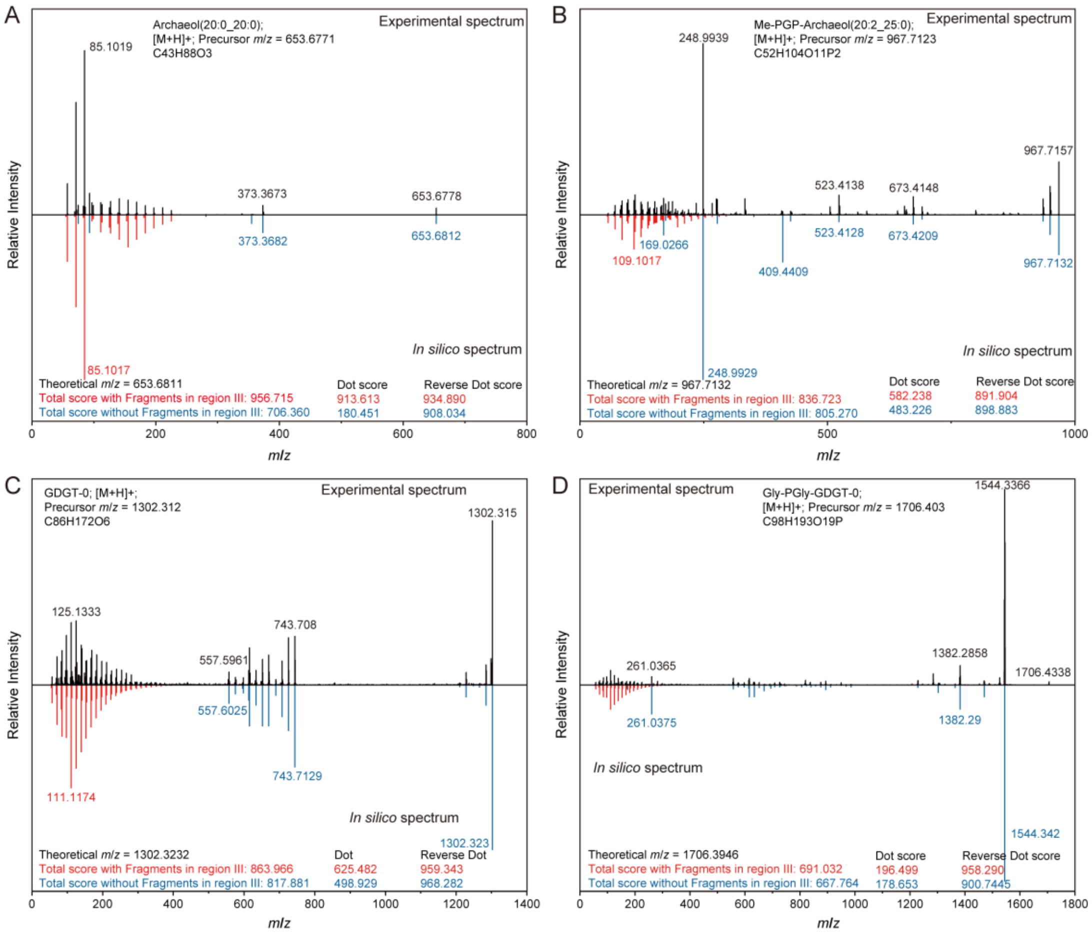

Figure S5. Effect of including low *m*/*z* fragment ions (Region III, m/z < 300) on spectral matching scores. A) Archaeol (20:0_20:0), $[M+H]^+$; B) Me-PGP-Archaeol (20:2_25:0), $[M+H]^+$; C) GDGT-0, $[M+H]^+$; D) Gly-(PGly)-GDGT-0, $[M+H]^+$. Dot score and reverse dot score were calculated using the MS-DIAL software. Fragments with (red) and without (blue) resulting from further cleavage of the carbon chains and polar headgroups. The experimental spectrum is shown at the top (black) and the *in silico* spectrum at the bottom (red and blue).

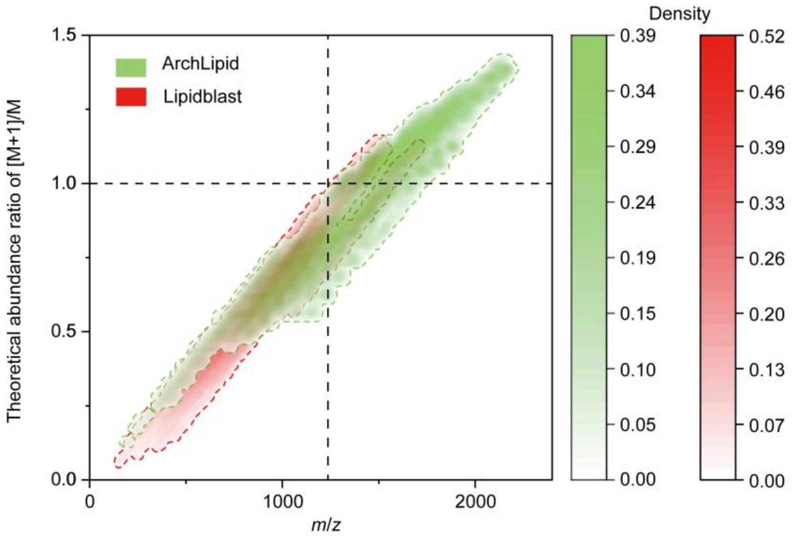


Figure S6. The theoretical abundance ratio of (M+1)/M was calculated based on the formula in the LipidBlast library and ArchLips database. Density refers to the distribution of individual lipids with the *m*/*z* values.

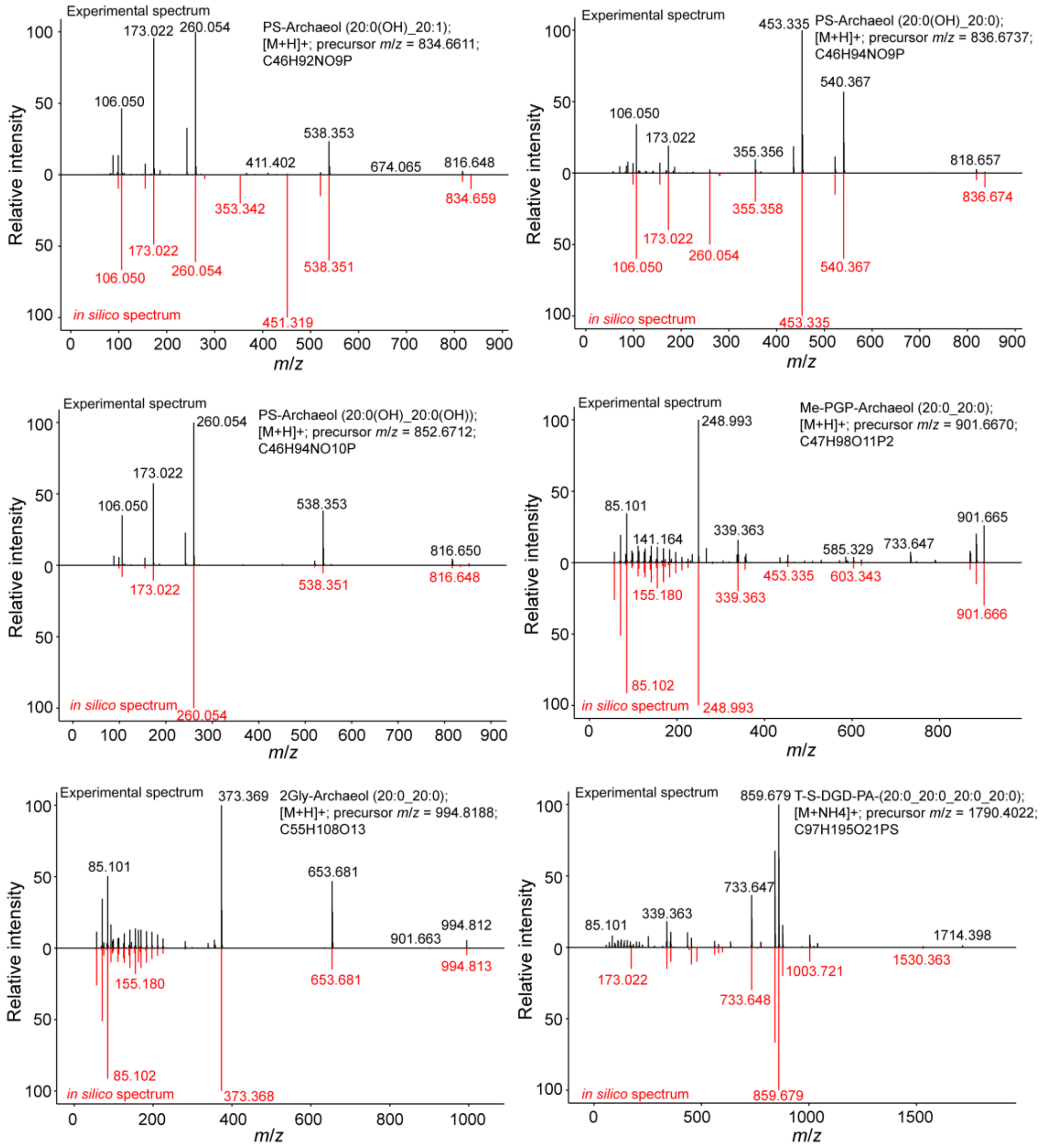


Figure S7. Examples of $MS^2$ library identification of archaeal ether lipids. The *in silico* spectrum in the library is in red (bottom), and the experimental spectrum is in black (top).

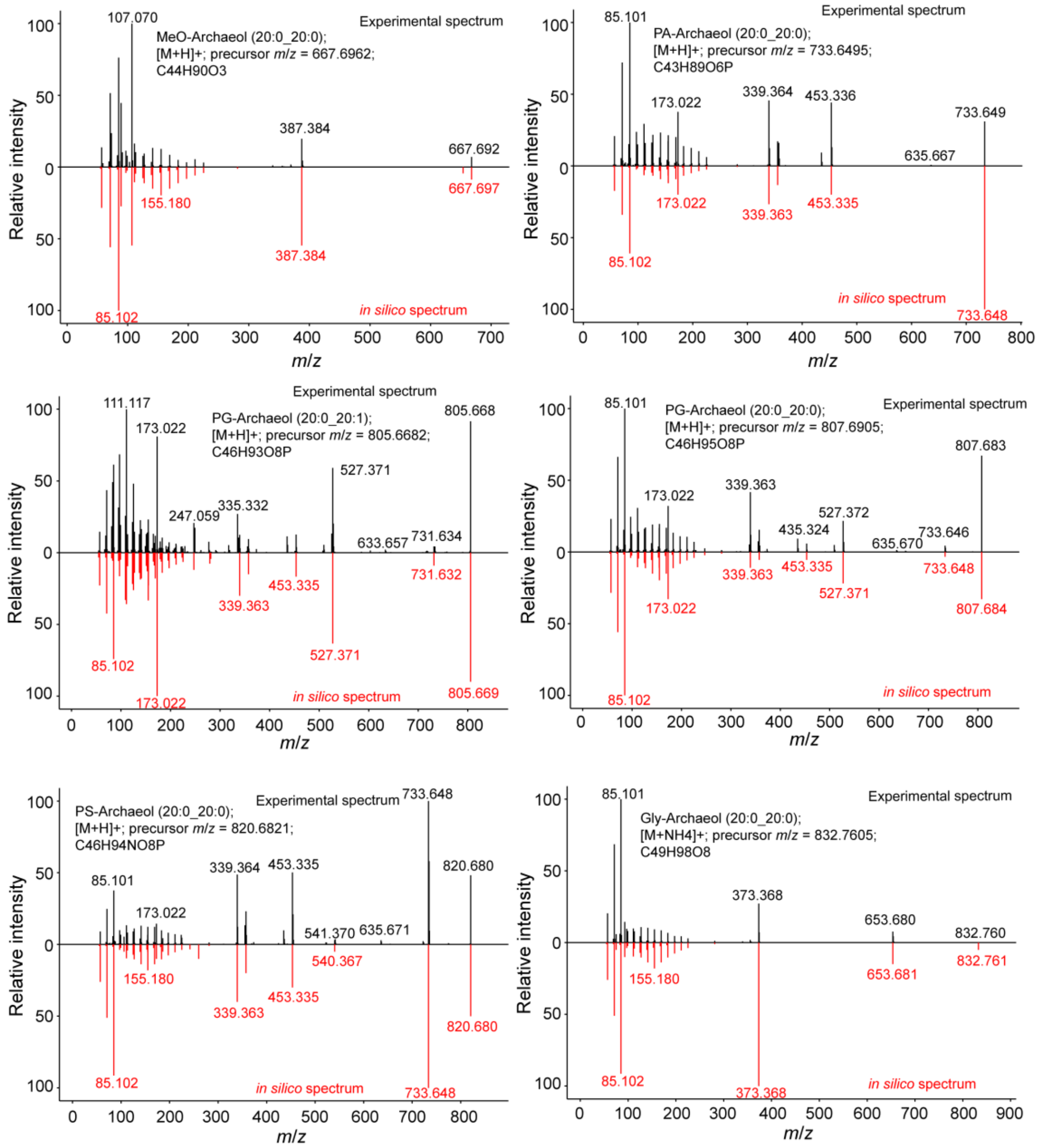


Figure S8. Examples of $MS^2$ library identification of archaeal ether lipids. The *in silico* spectrum in the library is in red (bottom), and the experimental spectrum is in black (top).

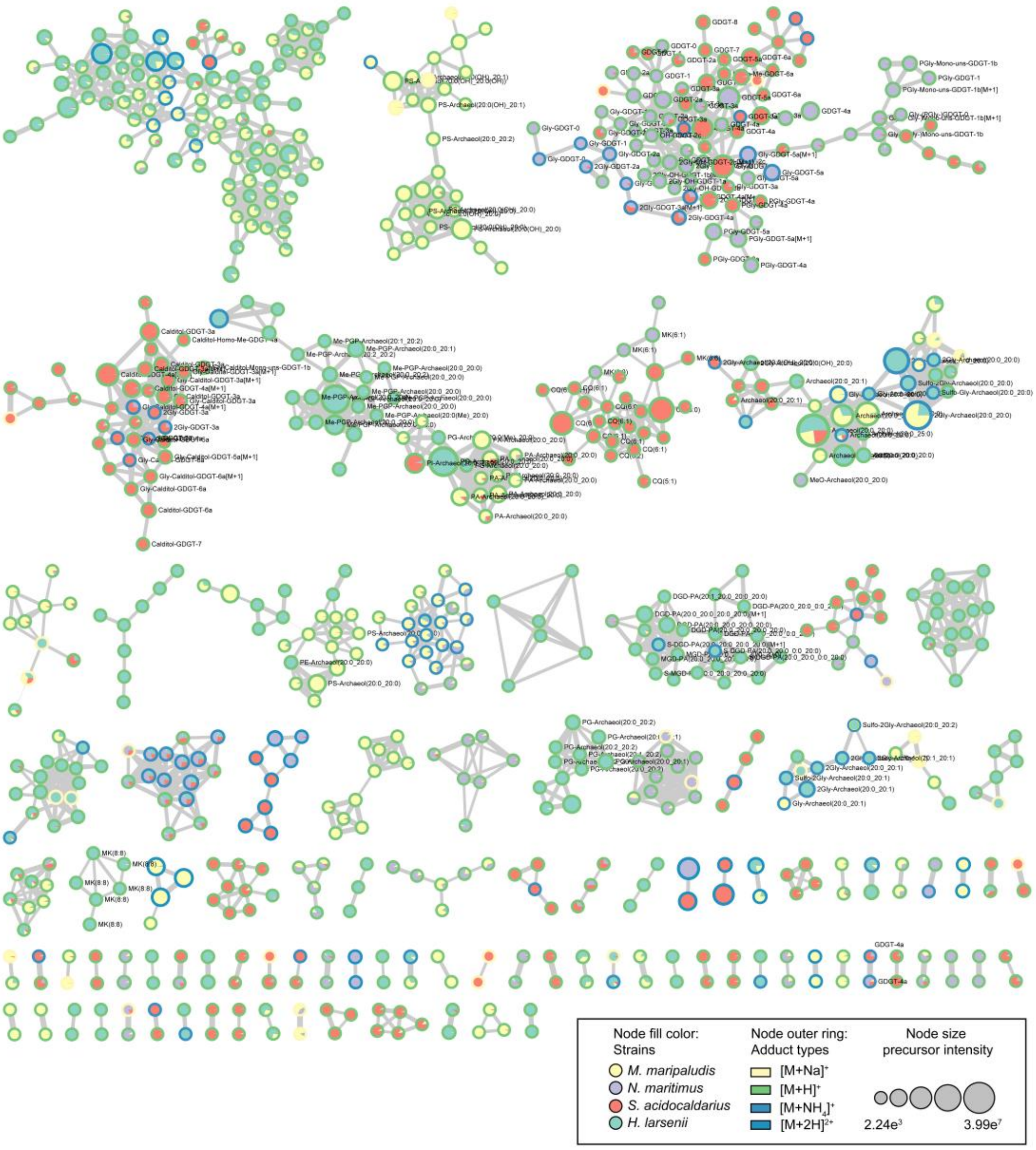


Figure S9. Feature-based molecular networking (FBMN) based on $MS^2$ spectra of lipid features of four representative archaeal strains. The singleton clusters with node <2 in FBMN were excluded. The filled color of the nodes indicates the distribution of lipid features of the four archaeal strains as shown in the legend, and the color of the outer rings refers to the adduct types. The width of the line connecting the nodes represents the level of mass spectral similarities, with thicker lines indicating greater similarity between the two nodes.

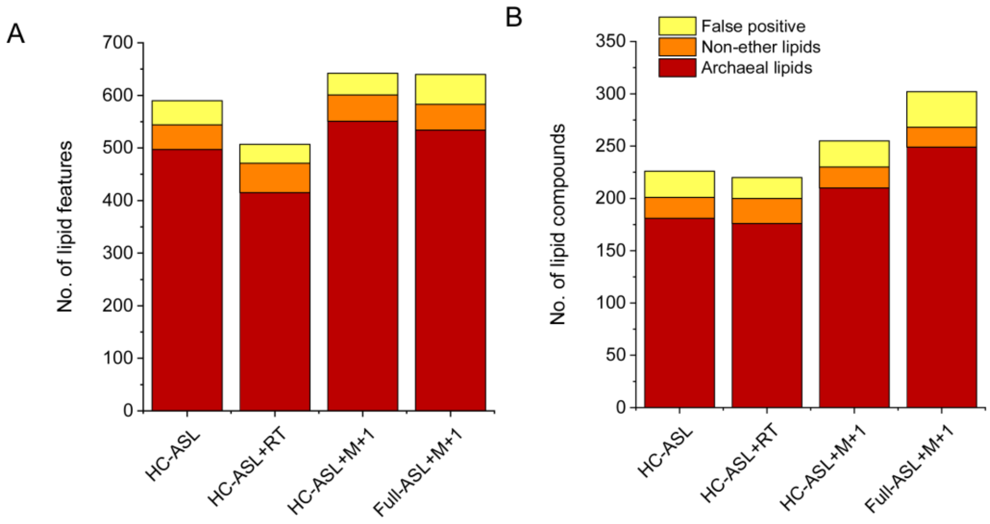


Figure S10. Number of annotated archaeal lipid features in environmental samples using different spectral libraries, with and without a retention time (RT) tolerance of 3 minutes. A) Number of lipid features annotated. B) Number of lipid compounds annotated.

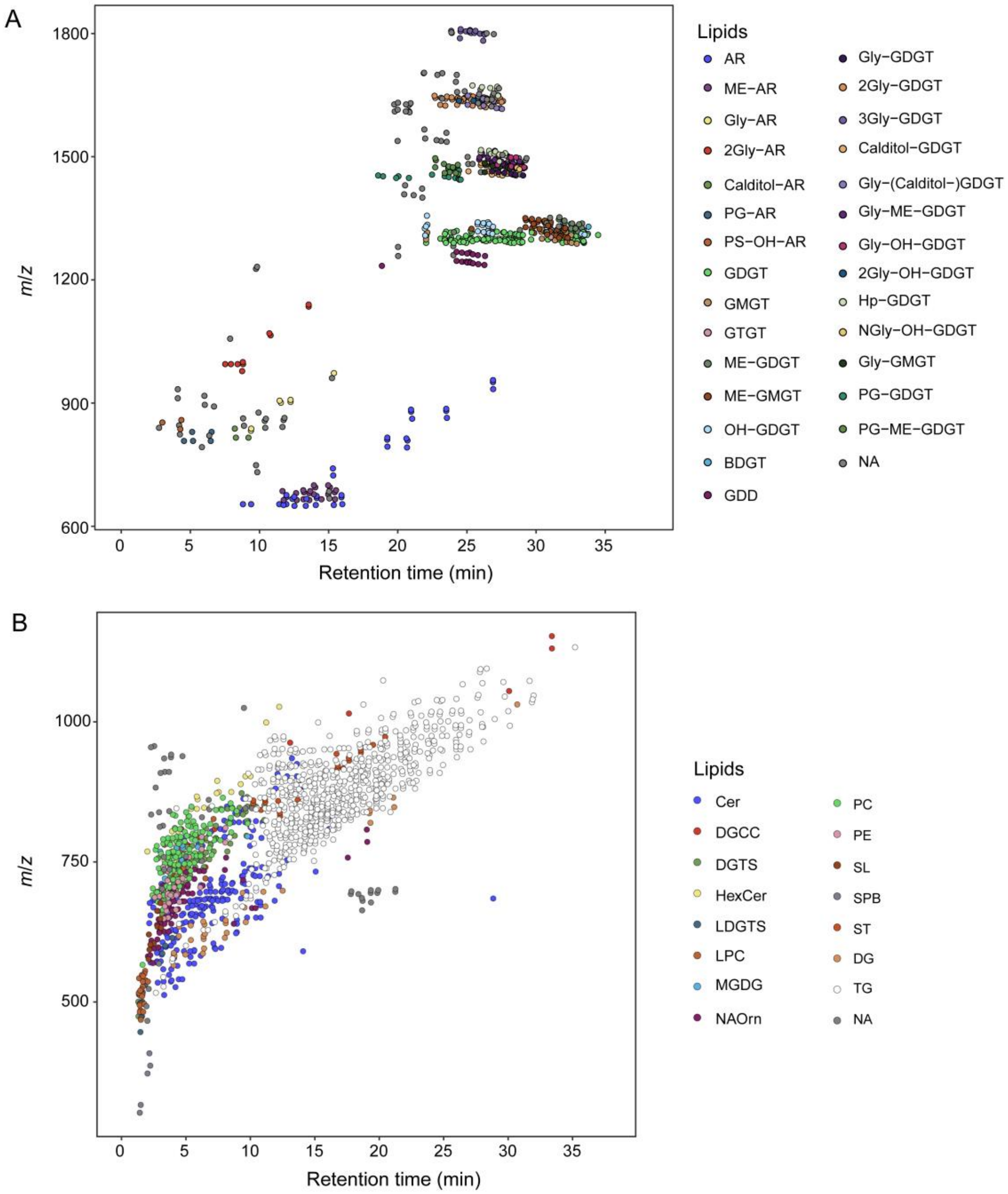


Figure S11 Retention time (RT) vs. *m*/*z* plot showing retention time and mass-to-charge ratio (*m*/*z*) of membrane lipids annotated with spectral matching of environmental samples. A) Archaeal lipids using the ArchLips database; B) bacterial-like lipids using the LipidBlast spectral library.

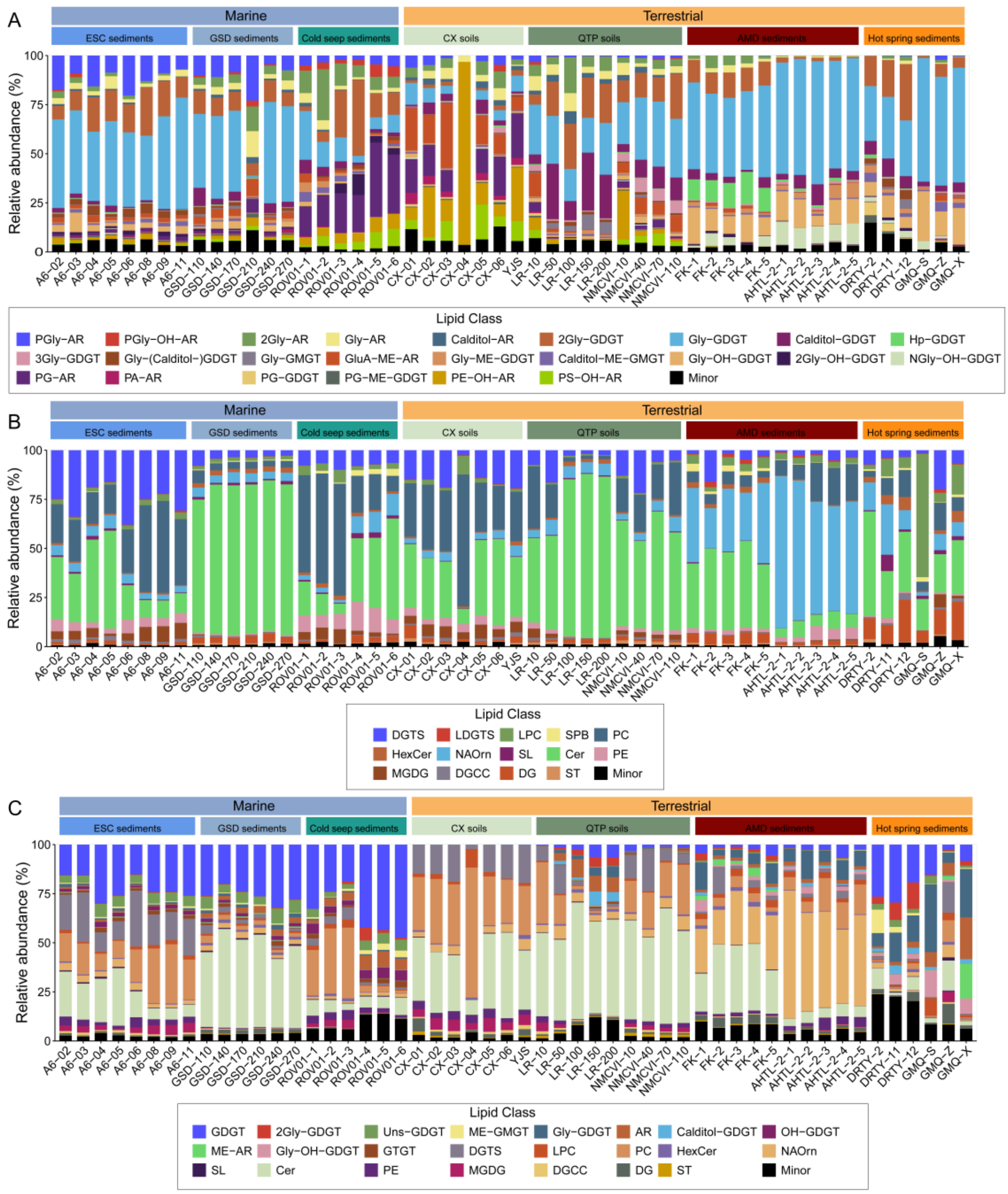


Figure S12. Relative abundance of archaeal and bacterial-like lipids identified using *in silico* mass spectral libraries across diverse environmental samples. A) Archaeal intact polar lipids (IPLs) identified using the ArchLips database. B) Bacterial-like lipids annotated with the LipidBlast library. C) Comprehensive environmental lipidomic profiles combining archaeal and bacterial-like membrane lipids.

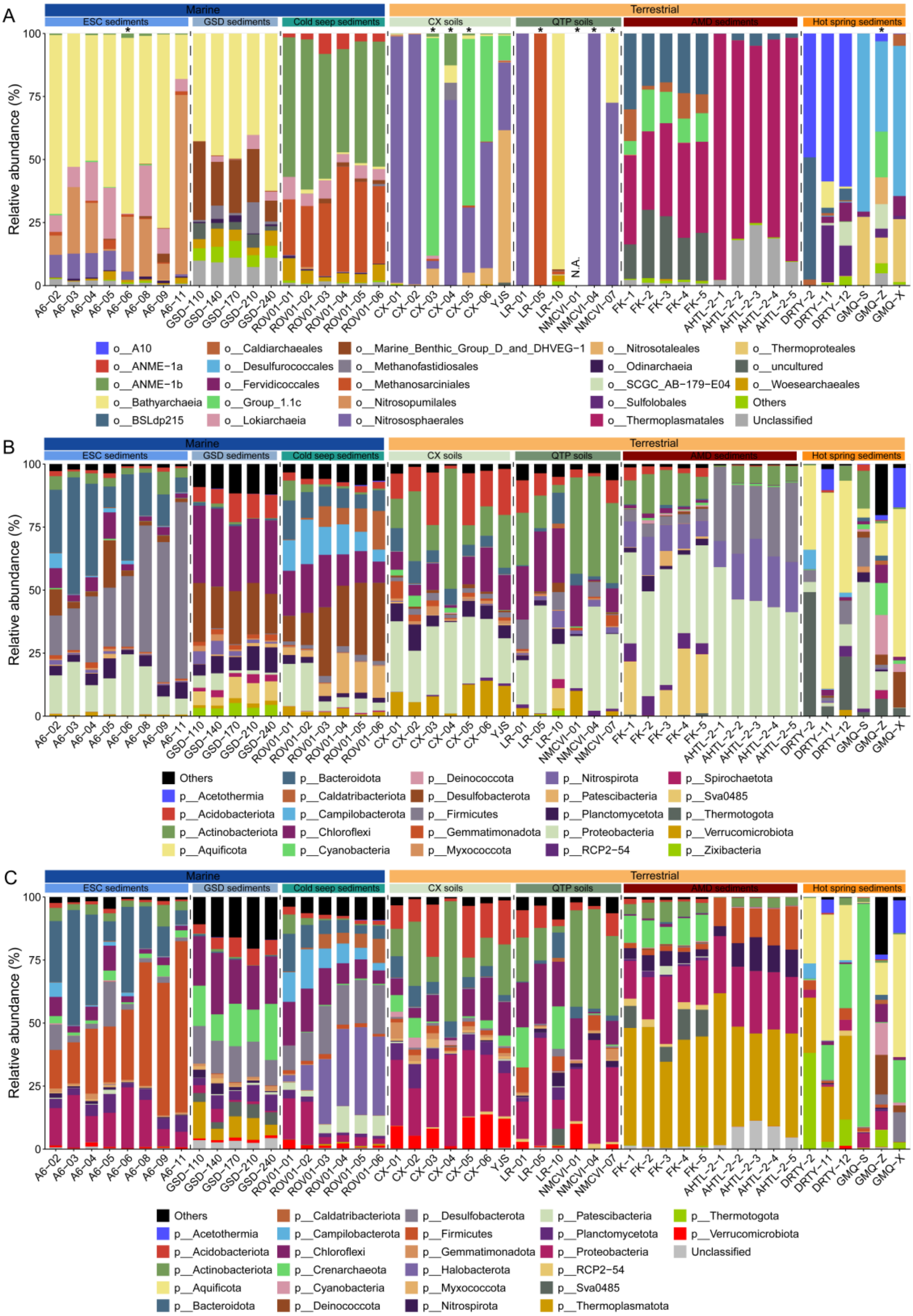


Figure S13. Relative abundance of archaeal and bacterial community identified using 16S rRNA sequencing with the 515F/806R primers across diverse environmental samples. A) Archaeal community composition. B) Bacterial community composition. C) Overall microbial community consisted of both archaeal and bacteria. An “*” indicates sample where archaeal read counts were fewer than 1,000.

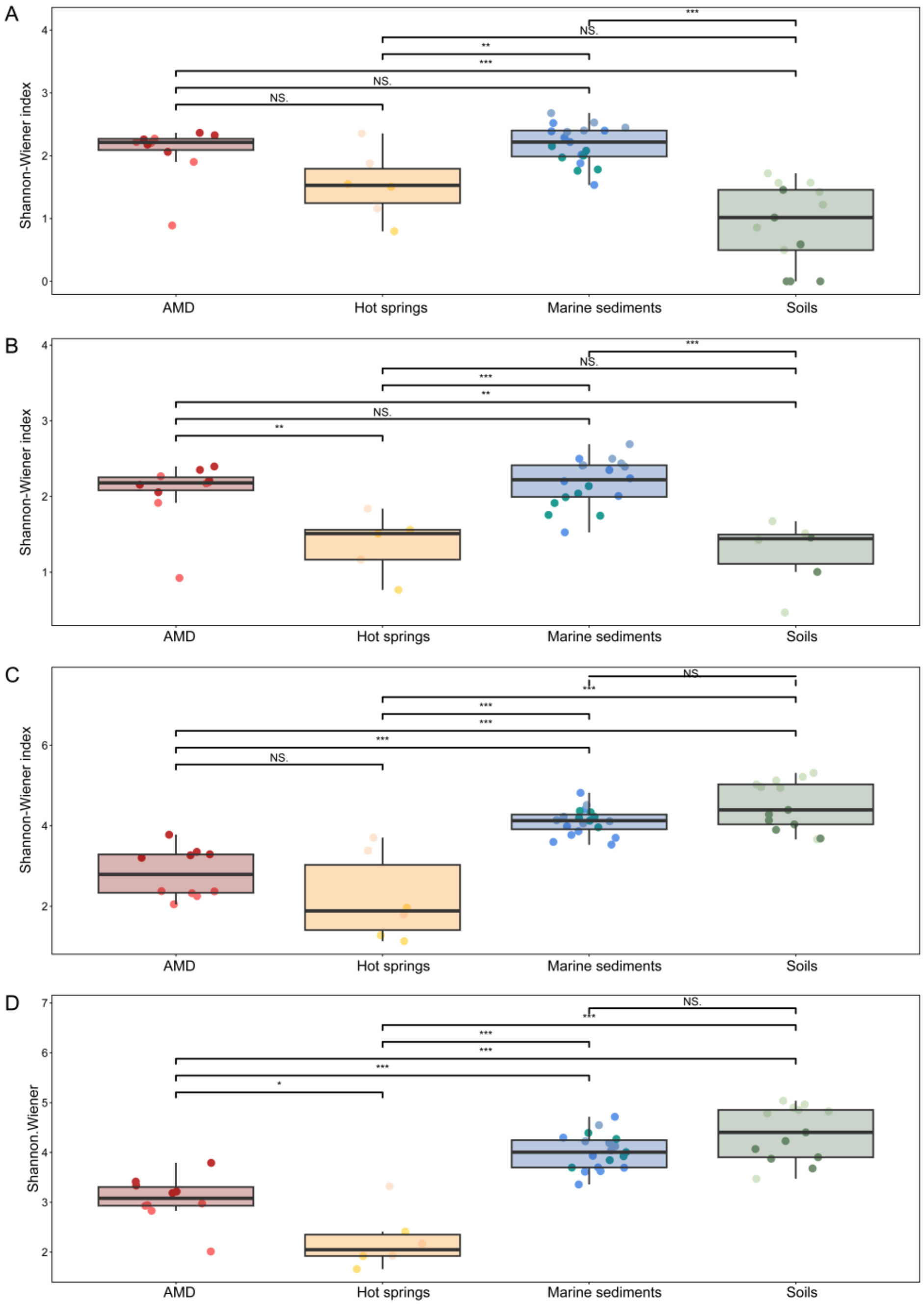


Figure S14 Microbial diversity of environmental samples. A) Shannon wiener index of archaeal community using all detected sequences. B) Shannon wiener index of archaeal community calculated after the archaeal sequences were normalized to 1,000. C Shannon wiener index of bacterial community. D) Overall microbial community includes archaeal and bacterial sequences. NS., not significant. The “*” indicates the significant level.

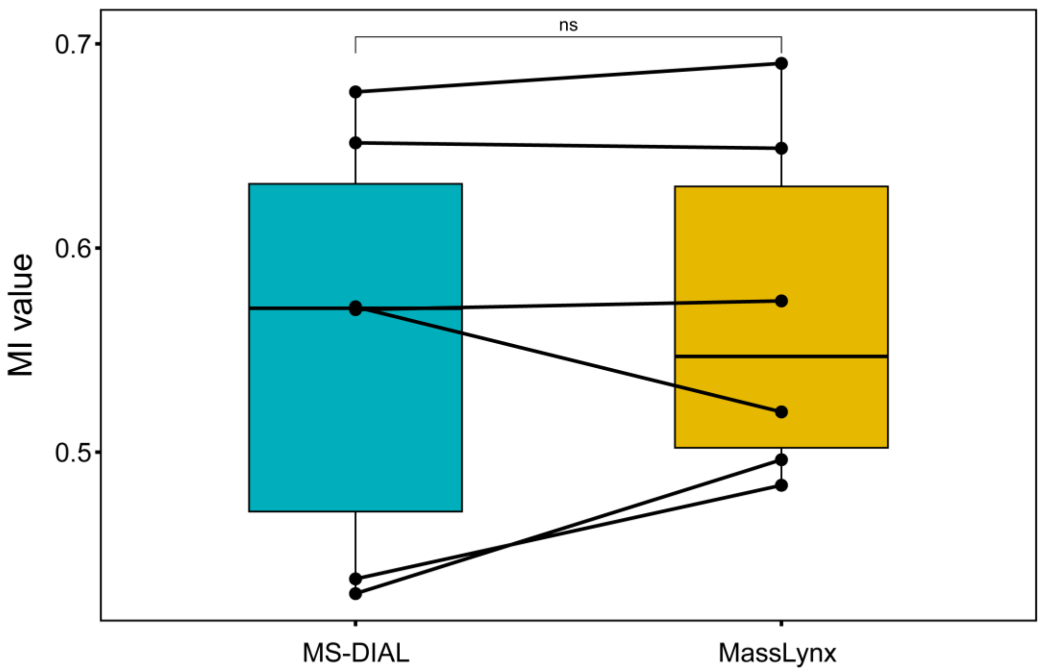


Figure S15. Boxplots comparing the Methane index (MI) calculated using the data automatically processed using MS-DIAL software and manual integration using the MassLynx software. A paired t-test was used with a significant level of $p < 0.05$. Three adducts of $[M+H]^+$, $[M+NH_4]^+$, and $[M+Na]^+$ were used, and the peak area was merged for proxy calculation.

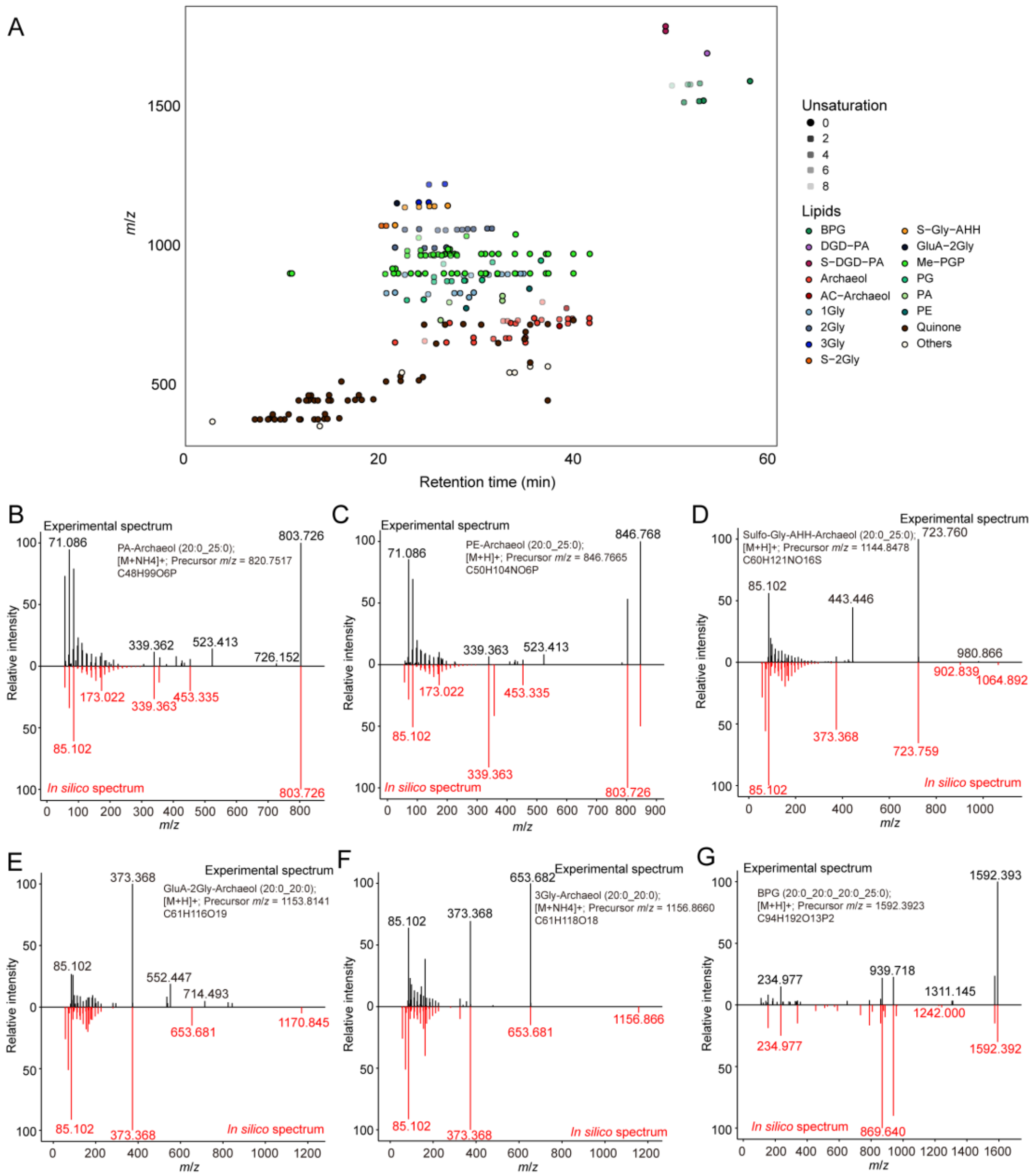


Figure S16. High-throughput lipid identification using the ArchLips database on a lipidomic dataset from four halobacterial strains analyzed with an Orbitrap MS platform[9, 10]. A) Retention time (RT) vs. *m/z* plot showing annotated lipid classes across the four halobacterial strains. Spectral matching of experimental MS² spectra (black, top) against referenced *in silico* spectra from the ArchLips database (red, bottom) for representative archaeal lipids.

Supplementary Tables 1-3

Table S1. Environmental sample information.

| Region | Sample Name | Sample types | Layers | Sampling date | Longitude (E) | Latitude (N) | Altitude[a]/Water depth (m) | pH | References |
|---|---|---|---|---|---|---|---|---|---|
| Western region of Sichuan | CX-01 | Soil | Surface | 2023.6 | 102.880 | 30.961 | 3867 [a] | 7.1 | This study |
| Western region of Sichuan | CX-02 | Soil | Surface | 2023.6 | 101.868 | 30.915 | 2409 [a] | 7.6 | This study |
| Western region of Sichuan | CX-03 | Soil | Surface | 2023.6 | 101.619 | 30.529 | 3541[a] | 5.9 | This study |
| Western region of Sichuan | CX-04 | Soil | Surface | 2023.6 | 101.540 | 30.431 | 3578 [a] | 8.1 | This study |
| Western region of Sichuan | CX-05 | Soil | Surface | 2023.6 | 101.641 | 30.183 | 3735 [a] | 6.4 | This study |
| Western region of Sichuan | CX-06 | Soil | Surface | 2023.6 | 101.588 | 30.211 | 4211 [a] | 5.9 | This study |
| Wuhan | YJS | Soil | Surface | 2016.9 | 114.435 | 30.514 | - | 5.1 | This study |
| Qinghai-Tibetan plateau | LR-01 | Soil | Subsurface | 2018.7 | 91.230 | 30.524 | 4243 [a] | 7.3 | Tang et al.,2023 |
| Qinghai-Tibetan plateau | LR-05 | Soil | Subsurface | 2018.7 | 91.230 | 30.524 | 4243 [a] | 6.3 | Tang et al.,2023 |
| Qinghai-Tibetan plateau | LR-10 | Soil | Subsurface | 2018.7 | 91.230 | 30.524 | 4243 [a] | 5.9 | Tang et al.,2023 |
| Qinghai-Tibetan plateau | NMCVI-01 | Soil | Subsurface | 2018.7 | 91.040 | 30.719 | 4845 [a] | 7.1 | Tang et al.,2023 |
| Qinghai-Tibetan plateau | NMCVI-04 | Soil | Subsurface | 2018.7 | 91.040 | 30.719 | 4845 [a] | 7.8 | Tang et al.,2023 |
| Qinghai-Tibetan plateau | NMCVI-07 | Soil | Subsurface | 2018.7 | 91.040 | 30.719 | 4845 [a] | 7.6 | Tang et al.,2023 |
| Tengchong | DRTY-2 | Hot spring | Surface | 2019.7 | 98.438 | 24.954 | - | 3.0 | This study |
| Tengchong | DRTY-11 | Hot spring | Surface | 2019.7 | 98.437 | 24.954 | - | 4.0 | This study |
| Tengchong | DRTY-12 | Hot spring | Surface | 2019.7 | 98.437 | 24.954 | - | 5.0 | This study |
| Tengchong | GMQ-S | Hot spring | Surface | 2019.7 | 98.436 | 24.951 | - | 9.0 | This study |
| Tengchong | GMQ-Z | Hot spring | Surface | 2019.7 | 98.436 | 24.951 | - | 9.0 | This study |
| Tengchong | GMQ-X | Hot spring | Surface | 2019.7 | 98.436 | 24.951 | - | 10.0 | This study |
| Fankou AMD | FK-1 | Acid mine drainage | Surface | 2017.9 | 113.663 | 25.049 | - | 3.1 | Hao et al., 2022 |
| Fankou AMD | FK-2 | Acid mine drainage | Surface | 2017.9 | 113.664 | 25.049 | - | 2.5 | Hao et al., 2022 |

| | | | | | | | | | |
|---|---|---|---|---|---|---|---|---|---|
| Fankou AMD | FK-3 | Acid mine drainage | Surface | 2017.9 | 113.664 | 25.049 | - | 5.5 | Hao et al., 2022 |
| Fankou AMD | FK-4 | Acid mine drainage | Surface | 2017.9 | 113.664 | 25.049 | - | 3.0 | Hao et al., 2022 |
| Fankou AMD | FK-5 | Acid mine drainage | Surface | 2017.9 | 113.663 | 25.050 | - | 2.6 | Hao et al., 2022 |
| Tongling AMD | AHTL-2-1 | Acid mine drainage | Surface | 2017.10 | 117.992 | 30.945 | - | 2.5 | Hao et al., 2022 |
| Tongling AMD | AHTL-2-2 | Acid mine drainage | Surface | 2017.10 | 117.993 | 30.945 | - | 2.6 | Hao et al., 2022 |
| Tongling AMD | AHTL-2-3 | Acid mine drainage | Surface | 2017.10 | 117.993 | 30.946 | - | 2.6 | Hao et al., 2022 |
| Tongling AMD | AHTL-2-4 | Acid mine drainage | Surface | 2017.10 | 117.992 | 30.946 | - | 2.6 | Hao et al., 2022 |
| Tongling AMD | AHTL-2-5 | Acid mine drainage | Surface | 2017.10 | 117.994 | 30.945 | - | 2.6 | Hao et al., 2022 |
| East China Sea | A6-02 | Marine | Surface | 2015.8 | 122.237 | 30.951 | 8 | N.A. | Chen et al., 2024 |
| East China Sea | A6-03 | Marine | Surface | 2015.8 | 122.382 | 30.909 | 12 | N.A. | Chen et al., 2024 |
| East China Sea | A6-04 | Marine | Surface | 2015.8 | 122.502 | 30.869 | 18 | 8.0 | Chen et al., 2024 |
| East China Sea | A6-05 | Marine | Surface | 2015.8 | 122.649 | 30.821 | 44 | N.A. | Chen et al., 2024 |
| East China Sea | A6-06 | Marine | Surface | 2015.8 | 122.807 | 30.773 | 30 | N.A. | Chen et al., 2024 |
| East China Sea | A6-08 | Marine | Surface | 2015.8 | 123.249 | 30.638 | 59 | 8.0 | Chen et al., 2024 |
| East China Sea | A6-09 | Marine | Surface | 2015.8 | 123.500 | 30.559 | 53 | N.A. | Chen et al., 2024 |
| East China Sea | A6-11 | Marine | Surface | 2015.8 | 124.000 | 30.408 | 45 | N.A. | Chen et al., 2024 |
| South China Sea | ROV01-01 | Cold seep | Subsurface | 2020.5 | 110.782 | 17.945 | 1736 | 7.7 | Zhang et al., 2023 |
| South China Sea | ROV01-02 | Cold seep | Subsurface | 2020.5 | 110.782 | 17.945 | 1736 | 7.7 | Zhang et al., 2023 |
| South China Sea | ROV01-03 | Cold seep | Subsurface | 2020.5 | 110.782 | 17.945 | 1736 | 7.9 | Zhang et al., 2023 |
| South China Sea | ROV01-04 | Cold seep | Subsurface | 2020.5 | 110.782 | 17.945 | 1736 | 7.9 | Zhang et al., 2023 |
| South China Sea | ROV01-05 | Cold seep | Subsurface | 2020.5 | 110.782 | 17.945 | 1736 | 7.9 | Zhang et al., 2023 |
| South China Sea | ROV01-06 | Cold seep | Subsurface | 2020.5 | 110.782 | 17.945 | 1736 | 7.9 | Zhang et al., 2023 |
| Pearl River | GSD-110 | Marine | Subsurface | 2017.10 | 113.806 | 22.132 | 21 | N.A. | Wang et al., 2020 |
| Pearl River | GSD-140 | Marine | Subsurface | 2017.10 | 113.806 | 22.132 | 21 | N.A. | Wang et al., 2020 |
| Pearl River | GSD-170 | Marine | Subsurface | 2017.10 | 113.806 | 22.132 | 21 | N.A. | Wang et al., 2020 |
| Pearl River | GSD-210 | Marine | Subsurface | 2017.10 | 113.806 | 22.132 | 21 | N.A. | Wang et al., 2020 |
| Pearl River | GSD-240 | Marine | Subsurface | 2017.10 | 113.806 | 22.132 | 21 | N.A. | Wang et al., 2020 |

Table S2. Head groups in the ArchLips database. Y = yes; N = no. Light blue color refers to glycolipids, green color phospholipids, and light yellow other compounds that cannot be classified into either glycolipids or phospholipids.

| Short name | Full Name | IPL-GDGT-IPL | Reported in diether lipids | Reported in tetraether lipids | Formula |
|---|---|---|---|---|---|
| Gly | Glycosyl | Y | Y | Y | C6H12O6 |
| 2Gly | Diglycosyl | Y | Y | Y | C12H22O11 |
| 3Gly | Triglycosyl | N | Y | Y | C18H32O16 |
| 4Gly | Tetraglycosyl | N | N | Y | C24H42O21 |
| Calditol | Calditol | Y | N | Y | C6H12O6 |
| Gly-Calditol | Glycosyl-calditol | N | N | Y | C12H22O11 |
| DeoxyGly | Deoxyglycosyl | N | Y | Y | C6H12O5 |
| G-2Gly | Glycerolglycosyl-glycosyl | Y | Y | N | C15H28O13 |
| G-Gly-NAcGly | Glycerolglycosyl-(N)-acetylglycosaminyl | N | N | N | C17H31NO13 |
| GluA | Glucuronic acid | Y | N | N | C6H10O7 |
| GluA-Gly | Glucuronic acid-glycosyl | N | N | N | C12H20O12 |
| GluA-2Gly | Glucuronic acid-diglycosyl | N | N | N | C18H30O17 |
| Hp | Heptosyl | N | N | N | C7H14O7 |
| Hp-Gly | Heptosyl-glycosyl | N | N | Y | C13H24O12 |

| | | | | | |
|---|---|---|---|---|---|
| MeOGly | Methoxylated glycosyl | N | N | N | C7H14O6 |
| MeOGly-Gly | Methoxylated glycosyl-glycosyl | N | Y | N | C13H24O11 |
| MeOGly-2Gly | Methoxylated glycosyl-diglycosyl | N | Y | N | C19H34O16 |
| NGly | Glycosaminyl | Y | N | N | C6H13NO5 |
| NAcGly | N-acetylglycosaminyl | Y | Y | N | C8H15NO6 |
| Ac-2Gly | Acetylglycosyl-glycosyl | N | Y | N | C14H24O12 |
| G-Gly-AcGly | Glycerolglycosyl-acetylglycosyl | N | Y | N | C17H30O14 |
| Sulfo-Gly | Sulphated glycosyl | N | N | N | C6H12O9S |
| Sulfo-2Gly | Sulfated diglycosyl | Y | Y | N | C12H22O14S |
| Sulfo-3Gly | Sulfated triglycosyl | N | Y | N | C18H32O19S |
| 2Sulfo-Gly | Disulfate-glycosyl | N | N | N | C6H12O12S2 |
| 2Sulfo-2Gly | Disulfate-diglycosyl | N | Y | N | C12H22O17S2 |
| Sulfo-Gly-AHH | Sulphated glycosyl-aminohexanehexaol | N | Y | N | C12H25NO14S |
| 2Sulfo-Gly-AHH | Disulphated glycosyl-aminohexanehexaol | N | Y | N | C12H25NO17S2 |
| SQ | Sulfoquinovosyl | N | N | N | C6H12O8S |
| SQ-Gly | Sulfoquinovosyl-glycosyl | N | N | N | C12H22O13S |
| SQ-2Gly | Sulfoquinovosyl-diglycosyl | N | N | N | C18H32O18S |

| | | | | | |
|---|---|---|---|---|---|
| Sulfo-SQ | Sulfated sulfoquinovosyl | N | N | N | $C_6H_{12}O_{11}S_2$ |
| SQ-Sulfo-Gly | Sulfoquinovosyl-sulphated glycosyl | N | N | N | $C_{12}H_{22}O_{16}S_2$ |
| SQ-AHH | Sulfoquinovosyl-aminohexanehexaol | N | N | N | $C_{12}H_{25}NO_{13}S$ |
| Sulfo-SQ-AHH | Sulphated sulfoquinovosyl-aminohexanehexaol | N | N | N | $C_{12}H_{25}NO_{16}S_2$ |
| Rib | Ribosyl | N | N | N | $C_5H_{10}O_5$ |
| NRib | N-ribosyl | N | N | N | $C_5H_{11}NO_4$ |
| Gly-Rib | Glycosyl-ribosyl | N | N | N | $C_{11}H_{20}O_{10}$ |
| Gly-PGly | Glycosyl-phosphatidylglycosyl | N | Y | N | $C_{12}H_{23}O_{14}P$ |
| Gly-PG | Phosphatidylglycerolhexose | Y | Y | N | $C_9H_{19}O_{11}P$ |
| Gly-PI | Glycosyl-phosphatidylinositol | N | Y | N | $C_{12}H_{23}O_{14}P$ |
| PE-PGly | Phosphatidylethanolamine-phosphatidylglycosyl | N | Y | N | $C_8H_{19}NO_{12}P_2$ |
| APT | Phosphoaminopentatetrol | N | Y | N | $C_5H_{14}NO_7P$ |
| N-Me-APT | N-monomethyl aminopentanetetrol | N | Y | N | $C_6H_{16}NO_7P$ |
| N-Me-APT-Me | N-monomethyl aminomethoxypentanetriol | N | Y | N | $C_7H_{18}NO_7P$ |
| N,N-2Me-APT | N,N-dimethyl aminopentanetetrol | N | Y | N | $C_7H_{18}NO_7P$ |
| N,N-2Me-APT-Me | N,N-dimethyl aminomethoxypentanetriol | N | Y | N | $C_8H_{20}NO_7P$ |
| N,N,N-3Me-APT | N,N,N-trimethyl aminopentanetetrol | N | Y | N | $C_8H_{20}NO_7P$ |

| | | | | | |
|---|---|---|---|---|---|
| N,N,N-3Me-APT-Me | N,N,N-trimethyl aminomethoxypentanetriol | N | Y | N | C9H22NO7P |
| NGly-PI | Glycosaminyl-phosphatidylinositol | N | N | N | C12H24NO13P |
| PA | Phosphatidic acid | Y | Y | N | H3O4P |
| 2Me-PA | Dimethyl-phosphatidyl acid | N | N | N | C2H7O4P |
| PC | Phosphatidylcholine | N | Y | N | C5H14NO4P |
| PE | Phosphatidylethanolamine | Y | Y | Y | C2H8NO4P |
| PME | Phosphatidyl-(N)-methylethanolamine | N | Y | Y | C3H10NO4P |
| PDME | Phosphatidyl-(N,N)-dimethylethanolamine | N | N | N | C4H12NO4P |
| PG | Phosphatidylglycerol | Y | Y | Y | C3H9O6P |
| PGly | Phosphatidylglycosyl | N | Y | Y | C6H13O9P |
| PGP | Phosphatidylglycerophosphate | N | Y | Y | C3H10O9P2 |
| Me-PGP | Methylated phosphatidylglycerophosphate | N | Y | N | C4H12O9P2 |
| PI | Phosphatidylinositol | Y | Y | Y | C6H13O9P |
| NAcGly-P | N-acetylglycosaminyl-phosphate | N | Y | N | C8H16NO9P |
| PPA | Pyrophosphate acid | N | N | N | H4O7P2 |
| Me-PPA | Methylated pyrophosphatidyl acid | N | N | N | CH6O7P2 |
| PS | Phosphatidylserine | Y | Y | N | C3H8NO6P |

| | | | | | |
|---|---|---|---|---|---|
| Gly-PS | Glycosyl-phosphatidylserine | N | Y | N | C9H18NO11P |
| Sulfo-PG | Phosphatidylglycerosulfate | N | Y | N | C3H9O9PS |
| NGly-PGly | Glycosaminyl-phosphatidylglycosyl | N | Y | N | C12H24NO13P |
| NAcGly-PGly | N-acetylglycosaminyl-phosphatidylglycosyl | N | Y | N | C14H26NO14P |
| 2NAc-2Gly-P | Di-(N)-acetylglycosaminyl glycosyl phosphate | N | Y | N | C16H29N2O14P |
| PIP | Phosphatidylinositolphosphate | N | N | N | C6H14O12P2 |
| PIP2 | Phosphatidylinositol-4,5-bisphosphate | N | N | N | C6H15O15P3 |
| PIP3 | Phosphoinositol-3,4,5-trisphosphate | N | N | N | C6H16O18P4 |
| PnC | Phosphonylcholine | N | N | N | C5H14NO3P |
| PnE | Phosphonylethanolamine | N | N | N | C2H8NO3P |
| CDP | Cytidine Diphosphate | N | Y | N | C9H15N3O11P2 |
| UDP | Uridine-diphosphate | N | N | N | C9H14N2O12P2 |
| MeO | Methoxylated | Y | Y | Y | CH4O |
| AC | Acetyl | N | Y | N | C2H4O2 |

Table S3. Annotation of archaeal lipids using the high confidence spectral library(HC-SL) and full spectral library (full-SL) of ArchLips database from LipidBlast and GNPS libraries that contain only bacterial-like lipids.

| Library | HC-SL | Full-SL |
|---|---|---|
| Lipidblast | 0.80% | 6.62% |
| GNPS | 0.19% | 0.60% |
